# Learning evolutionary parameters from genealogies using allelic trees

**DOI:** 10.1101/2025.03.03.641273

**Authors:** Antoine Aragon, Amaury Lambert, Thierry Mora, Aleksandra M. Walczak

## Abstract

Cellular diversification in processes from development to cancer progression and affinity maturation is often linked to the appearance of new mutations, generating genetic heterogeneity. Describing the underlying coupled genetic and growth processes that result in the observed diversity in cell populations is informative about the timing, drivers and outcomes of cell fates. Current approaches based on phylogenetic methods do not cover the entire range of evolutionary rates, often making artificial assumptions about the timing of events. We introduce CBA, a probabilistic method that infers the division, degradation and mutation rates from the observed genetic diversity in a population of cells. It uses a summarized backbone tree, intermediary between the true cell tree and the allelic tree representing the ancestral relationships between types, called a monogram, which allows for efficient sampling of possible phylogenies consistent with the observed mutational signatures. We demonstrate the accuracy of our method on simulated data and compare its performance to standard phylogenetic approaches.

## I. INTRODUCTION

The growth and diversification of somatic cell populations within a organism offer a perfect example of asexual evolution. Particular instances include cancer progression [1–3], affinity maturation of antibodies [4–10], and haematopoiesis [11–14]. Because these processes may be modeled by traditional evolutionary genetics [15], inference methods developed in the fields of phylogenetics and population genetics could be adapted to estimate the evolutionary parameters of somatic processes, such as division, death, and mutation rates, as well as their dependence on time, tissue or cell type. Efforts in adapting such methods to somatic evolution hold the potential to gain new insight into a variety of biological processes, such as the timing and drivers of cell fate decision during early development [16–18], the mechanisms of hyper-mutation and selection during antibody affinity maturation [6–10], or the drivers of cancer onset and progression [19].

Several methods exist to infer the phylodynamics of growing populations. Bayesian inference is widely used to infer phylogenetic trees from molecular sequences and to test evolutionary hypotheses. In particular, the “Swiss knife” BEAST 2.0 platform [20], based on Monte-Carlo Markov chains, offers a large diversity of evolutionary models allowing for various inference tasks. Other methods have been proposed to address specific aims more efficiently. Starting from time-resolved genealogies, phylodyn allows for inference of time-dependent population size [21]. A distinct method, cloneRate, is optimized to efficiently infer the net growth rate [22], with applications to haematopoiesis. Starting from densely sampled sequence data of rapidly evolving populations such as viruses, TreeTime [23] infers time-resolved genealogies as well as evolutionary models, while another method infers their individual fitnesses [24]. Recently, a method was proposed to infer the mutation rate and the death-to-division ratio from phylogenetic trees with no time information, assuming that mutations occur at cell division [25].

A major challenge faced by inference methods is that we never have access to the full process that produced the observed data. Typical datasets will only include a sample of existing types, which are defined by their somatic DNA mutations, barcoding, or gene expression profiles measured by RNAseq (Fig. 1a). Estimating evolutionary parameters can be done directly from the whole cell lineage tree (Fig. 1b, left). However, this is not possible since cell deaths leave no signatures in the data. Phylogenetic approaches often start from the “reconstructed” tree, which traces the time-resolved genealogy of extant cells (Fig. 1b, second left), or from the reconstructed sub-tree of sampled cells (Fig. 1b, second right) called chronogram. However, what the data provides is a mixture of variants, or “types”, that arose at different, unknown times. Its phylogeny may be inferred from data to get a genealogical tree of extant cells whose branch lengths represent genetic distance, called a phylogram. When the abundance of each type may not be well measured, as we will assume in this paper, the phylogram boils down to a “tree of mutations,” called allelic tree (Fig. 1b, right), where each tip represents a type present in the sample, and each branch represents a series of mutations necessary to transition from one type to another.

**FIG 1:**
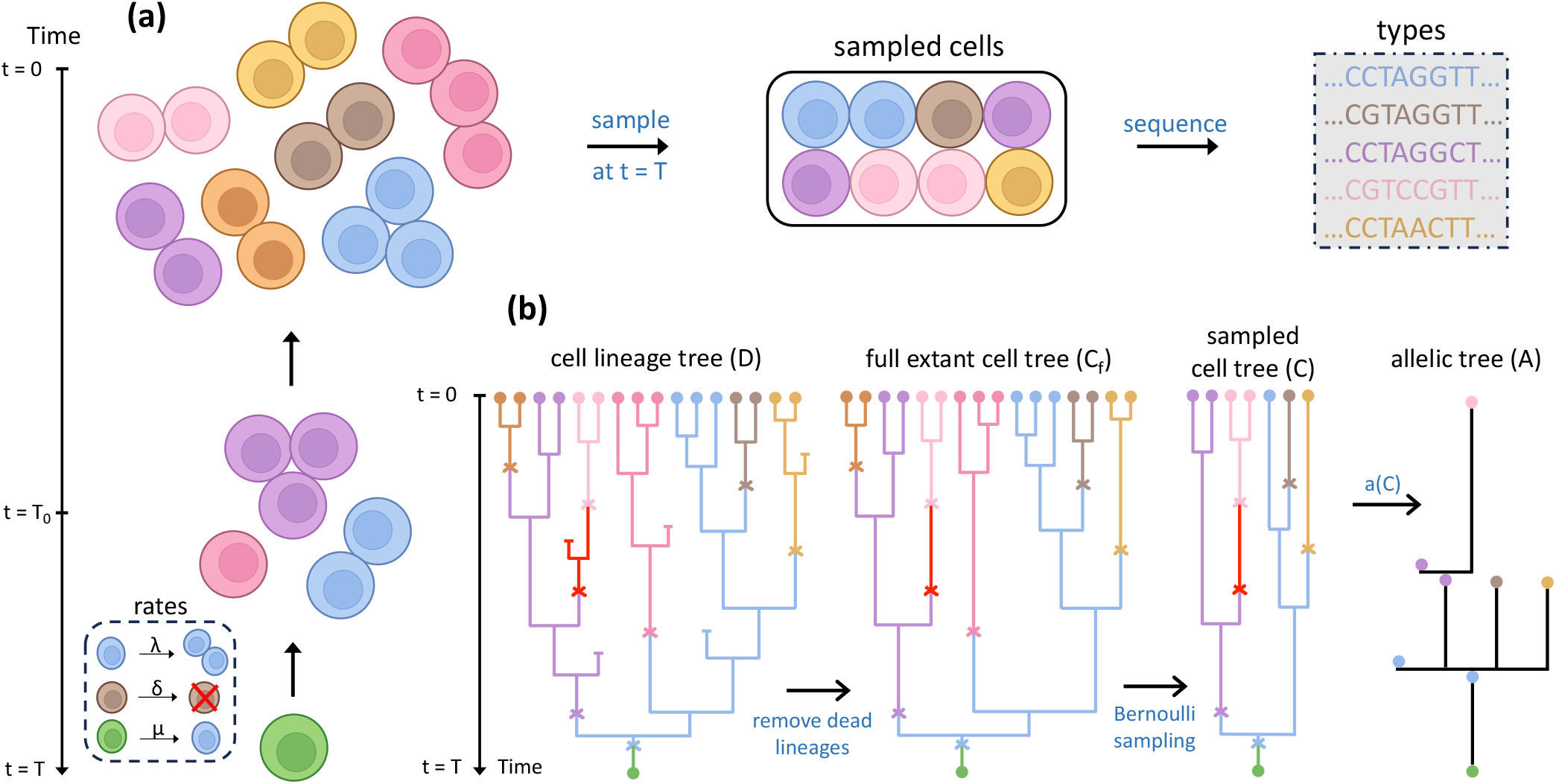
Evolutionary model of growing populations of cells. (a) Growing population of cells along with schematic of how data is obtained from typical experiments. The population is sampled at time *t* = 0 and then sequenced, yielding a list of (geno)types. (b) Left and middle-left panels: cell lineage tree and extant cell lineage tree generated by the population the population generates. Cells divide at rate *λ*, die at rate *δ* and acquire mutations at rate *µ*. In the cell tree *D*, cell deaths are depicted by a horizontal line marking the end of a lineage, and mutations are indicated by crosses. Each mutation corresponds to a change of type, which is represented by a change in color (infinite-allele model). The full extant cell lineage tree (*C*_*f*_) is the cell lineage tree where dead linages are removed. By contrast the cell tree *D* also represents dead cells. Middle-right panel: the sampled cell tree *C* is obtained through a Bernoulli sampling with parameter *ρ* on the leaves of *C*_*f*_. Right panel: The allelic tree *A* = *a*(*C*), obtained by keeping only the list and order of mutations across types. The length of branches in *A* indicates the number of mutations separating the types.

Many methods approximate the chronogram by the phylogram, by using the genetic distance between two cells as a rough measure of calendar time. That approximation is valid when the genetic distance is neither too small (to avoid small number fluctuations) nor too large, and under the double assumption that mutations are neutral and of constant rate—the molecular clock hypothesis. The genetic divergence between two individuals is on average 2*µℓt*, where *t* is the time since their most recent common ancestor, *µ* is the per nucleotide mutation rate and *ℓ* is the number of nucleotides in the stretch of DNA considered. In phylogenetics, individuals are usually sampled in different species so that *t* is large. In viral evolution [26], *t* may be small but *µℓ* is large.

In contrast, in somatic evolution *µℓt* may be small, so that the phylogram is a poor proxy of the chronogram. In specific cases, mutations can be artificially induced through genome editing-based lineage tracing methods to get a better resolution of time and apply classical methods with some adaptations [27–31]. In many cases however, inferring the chronogram from data is not possible. On the other hand, since mutations are rare, it is relatively straightforward to build the phylogram by maximum parsimony [4].

Our goal is to propose a probabilistic method that infers the division, degradation and mutation rates from the phylogram, which works even in the low mutation limit. Our strategy is based on an intermediate object between the chronogram (cell lineage tree) and the phylogram (allelic tree): the *monogram*, which we will also call backbone tree. The monogram can be viewed as a phylogram for which times of division and mutation events have been resolved and calibrated in time. The method we introduce, called CBA, interpolates between these three levels of description: (C) cell tree or chronogram, (B) backbone tree or monogram, and (A) allelic tree or phylogram.

## II. RESULTS

### A. Outline of the approach

We consider a population of cells that proliferates and mutates following a birth-death process with division rate *λ*, death rate *δ* and a neutral mutation rate *µ* (Fig. 1a). We assume that mutations accumulate with absolute time and not upon division. Time is defined in a backwards manner, such that *t* = *T* marks the beginning of the process while *t* = 0 marks the present. As the population grows, it generates a cell lineage tree *D* shown in Fig. 1b, which tracks divisions, deaths and mutations. The cell lineage tree is what an ideal observer would see given the evolution of the population between *t* = *T* and *t* = 0, as it contains all the past relevant information about growth and diversification. Finding estimates for the rates from *D* is easily accomplished through maximum likelihood using the branching property [32]. Unfortunately, in most datasets we do not have access to the whole tree but only have access to a sample of cells at one timepoint, or a few timepoints.

Removing death events that give rise to dead end branches from the cell tree *D* in Fig. 1b gives the lineage tree *C*_*f*_ of extant cells at *t* = 0. This object is often studied in macroevolution [33, 34]. A quasi ideal observer, seeing a population of living cells at time *t* = 0 with all their genetic distances, could only reconstruct *C*_*f*_ and not *D* since the present population contains no information about past death events. The likelihood of the reconstructed cell lineage tree was derived in [32] and could be used to infer the rates using maximum likelihood (see SI Appendix 1).

Typical experiments give access to neither *D* nor *C*_*f*_. The population is sampled at time *t* = 0 and then sequenced, yielding a list of (geno)types, in which abundances (number of cells representing each type) are often discarded since experimental biases (sequence amplification error, expression noise) make them unreliable. We call *C* the subtree of *C*_*f*_ that contains only the phylogeny of sampled cells, obtained by Bernoulli sampling on the leaves of *C*_*f*_ (Fig. 1b, second left). But experiments do not give direct information about the times of bifurcation and mutation events in *C*, nor about tree bifurcations that are not followed by mutations, and so in practice even the sampled tree *C* is unknown. Instead, experiments typically give the allelic tree *A*, which describes the history of mutations leading to the sequences. The allelic tree may be inferred from the list of type sequences using well established maximum likelihood methods such as IQ-Tree [35] and RAxML [36]. *A* = *a*(*C*) can be deduced from *C* (Fig. 1b), where the function *a* consists in removing branches in *C* with no mutations and making the length of all remaining branches proportional to the number of mutations. No explicit expression for the likelihood *P* (*A*| *λ, δ, µ*) exists for the allelic tree, making inference challenging.

Our method fills this gap and infers the rates *λ, δ, µ* governing the dynamics of the process through the allelic tree. The lack of an explicit expression for the likelihood of *A* makes this a difficult task. To overcome this hurdle, we connect *A* to *C* through the backbone tree *B* (Fig. 2). The backbone tree is an intermediate object between *A* and *C*. It is a function *B* = *b*(*C*) of the extant cell lineage tree *C*. The rules of *b* for transforming *C* into *B* are illustrated in Fig. 2. Subtrees downstream of terminal mutations, i.e. subtrees that have no further mutations (brown, yellow, and pink), are replaced by a single terminal branch. Subtrees downstream of internal (non-terminal) mutations (blue, purple, and red) are collapsed to only keep the part of the tree leading to further mutations. Mutations are boxed if in *C* there exists a path between the mutation and the present with no further mutation, in other words, if the type that emerged following that mutation is represented in the data. Mutations are circled if there is no such path and all sampled descendants carry additional mutations (e.g. the red mutation). The allelic tree *A* = *f* (*B*) can then be obtained from the backbone *B* through a function *f* which removes any information about the times of division and mutations, removes unrepresented types, and makes branch lengths proportional to the number of mutations (Fig. 2). It follows that *a*(*C*) = *f* (*b*(*C*)).

**FIG 2:**
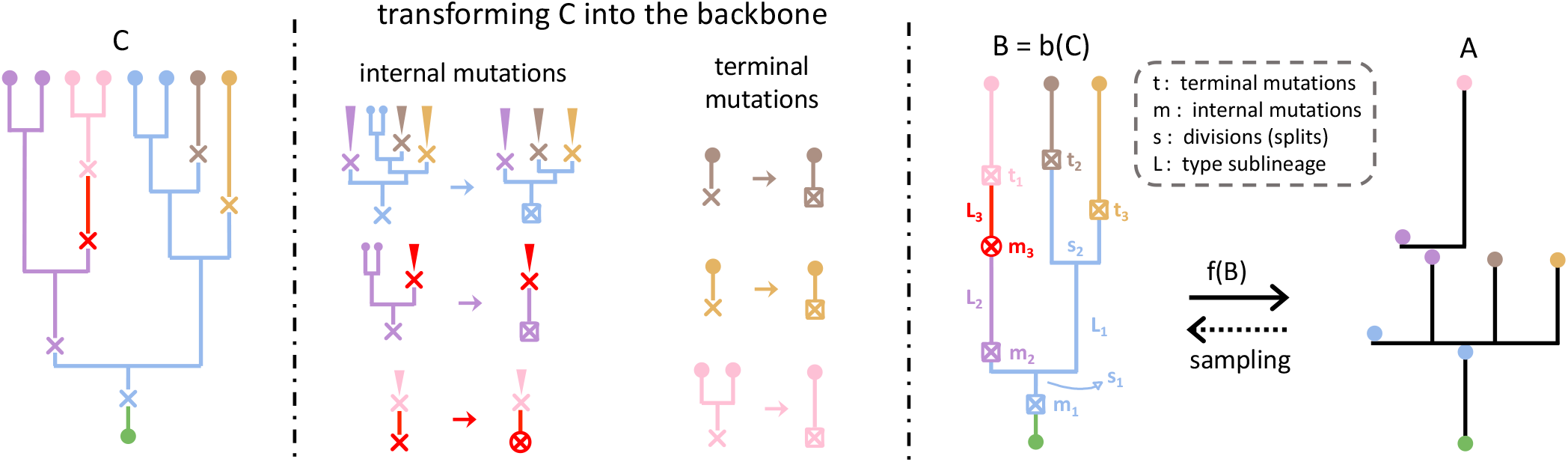
The backbone tree as an intermediate between cell and allelic trees. Left panel: the starting point is the tree of sampled extant cells *C*. Middle panel: set of rules transforming a cell tree into a backbone tree. Subtrees descending from terminal mutations (pink, brown, yellow) are collapsed into a single edge and the mutation is boxed to indicate that the type emerging from that mutation is observed. Internal mutations leading to subtrees carrying no further mutations are boxed, and the corresponding unmutated subtrees removed (blue, purple). Internal mutations from which no such subtree stems (red) are circled. Right panel: The backbone resulting from *B* is transformed into an allelic tree through *f* (*B*), so that *f* (*b*(*C*)) = *a*(*C*).

**FIG 3:**
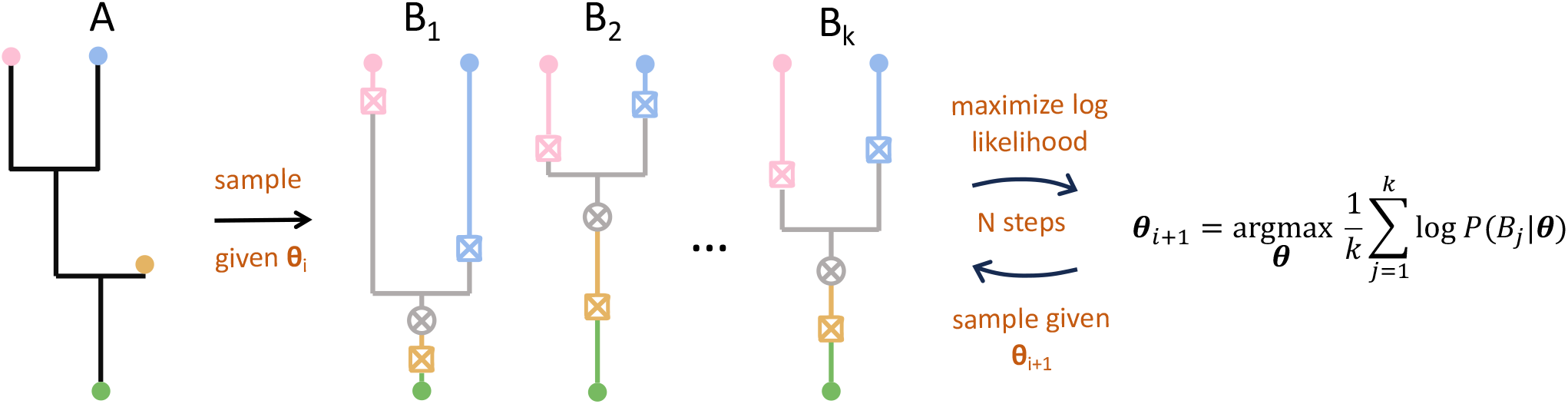
Expectation-maximization. Schematic of the inference scheme based on expectation maximization for one allelic tree *N* = 1. Given the current parameter estimates ***θ***_*i*_, backbone trees *{B*_1_, *B*_2_, …, *B*_*k*_*}* are sampled from *P* (*B A*, |***θ***_*i*_) using a Monte-Carlo Markov chain. Their mean log likelihood (expectation) is maximized (maximization) with respect to ***θ***, yielding the next parameter estimates ***θ***_*i*+1_. These expectation and maximization steps are repeated a total of *N*_MC_ times.

The backbone tree *B* is the simplest reduction of the cell tree *C* towards *A* that we found and whose likelihood *P* (*B*| *λ, δ, ν*) can still be computed analytically. It is close enough to the allelic tree so that fast sampling schemes exist to generate random backbone trees consistent with a given allelic tree, *B* ∈ *f* ^−1^(*A*). We exploit this last point to develop an efficient inference procedure.

### B. Likelihood of the backbone tree

To implement our approach we must first derive the likelihood of the backbone tree. A full derivation with explicit calculations is detailed in SI Appendix 2. This likelihood includes terms corresponding to divisions (splits) at times (*s*_1_, …, *s*_*n*−1_), internal boxed mutations at times (*m*_1_, …, *m*_′_), internal circled mutations at times (*m*_′ +1_, …, *m*), and terminal mutations at times (*t*_1_, …, *t*_*n*_), where *ℓ* is the total number of mutations, and *ℓ*^′^ the number of those which are boxed. All those times take values between 0 and *T*. The concatenation of intervals downstream of an internal mutation *i* until the next mutations is denoted by *L*_*i*_ and represented by the same color as the mutation in the left panel of Fig. 2. Putting all the division and mutation terms together, the likelihood of the backbone tree can be written as

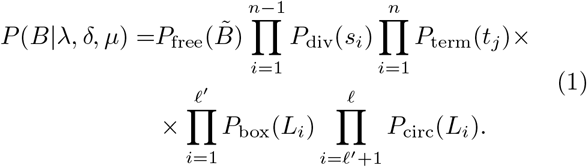

For example, for the tree in Fig. 2 the likelihood is:

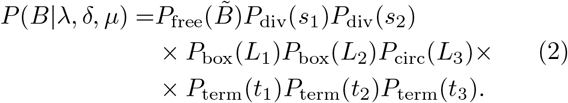

Each term is detailed below. While all mutations occur with rate *µ* and divisions with rate *λ*, in the backbone tree these rates must be corrected to account for the fate of the downstream subtrees, in particular that they must be represented in the data, and whether they undergo further mutations or not. To calculate the functional transforming C into the backbone form of the terms in the likelihood, it is useful to denote by *p*(*t*) the probability that a sublineage downstream of a point on the tree at time *t* survives until the present, and by *q*(*t*) the probability that that sublineage survives and also mutates at least once. These probabilities can be computed analytically, as shown in SI Appendix 2.

By construction, the backbone tree only carries a visible division event if both post-division branches mutate after the division. The rate of a division occurring at time *t* in *B* (Fig. S2) is then proportional to the division rate *λ* times the probability that each sublineage survives and mutates:

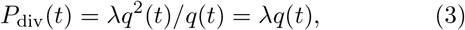

where the *q*(*t*) in the denominator comes from conditioning on the pre-division branch having at least one mutation and surviving.

The rate of a terminal mutation occurring at time *t* is given by the mutation rate *µ* times the probability *p*(*t*) − *q*(*t*) that the branch survives until the present but without any further mutation (Fig. S2), conditional again on the the pre-division branch surviving with at least one further mutation:

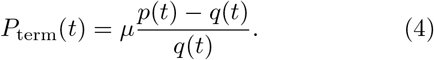

We see from both the *P*_div_(*t*) and *P*_term_(*m*_*j*_) terms that conditioning on the pre-division branch surviving introduces a non-obvious factor to the rates.

An internal circled mutation, such as the red mutation at time *m*_3_ in Fig. 2, means that there was no division event between this mutation and the next downstream mutations (in this case during the interval *L*_3_) that would lead to a sublineage surviving until the present with no further mutations — the existence of such a sublineage would imply the representation of the type emerging upon that mutation, making it a boxed mutation. The rate of such division events is given by 2*λ*(*p*(*t*) − *q*(*t*)), where the factor of 2 stems from the fact that there are 2 potential subtrees at each division that could cause the type to be represented (Fig S2). More rigorously, a division event can lead to 3 alternatives: two surviving subtrees each with at least one mutation with probability *q*(*t*)^2^, both without a mutation in the surviving branches with probability (*p*(*t*) − *q*(*t*))^2^, and one of each type with probability 2 *λ q*(*t*)(*p*(*t*) − *q*(*t*)) (Fig. S2). We must condition on the pre-division branch surviving with at least one mutation, since we are considering points on the tree between internal and terminal mutations. This means keeping only the case where there is a subtree of each type, since we want the type to be represented, and dividing its probability by *q*(*t*): 2*q*(*t*)(*p*(*t*)−*q*(*t*))*/q*(*t*) = 2(*p*(*t*)−*q*(*t*)).

As a result, the rate of mutations *not* followed by such division events until the next mutations is:

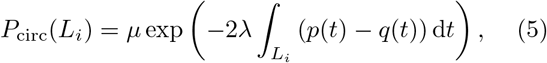

Internal boxed mutations occur with complementary rate, so that:

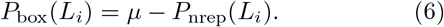

Finally, 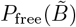 corresponds to the probability that no other event occurs on the backbone tree upstream of the terminal mutations (denoted by 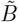):

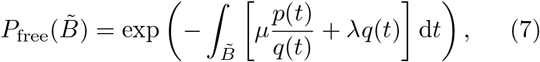

where the integral runs over all branches of 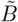. The first term describes the rate of mutations leading to surviving sublineages, conditioned on the pre-division branch surviving and having at least one further mutation. The second term corresponds to the rate of divisions (3).

### C. Inference procedure

Starting from one or multiple allelic trees *A* generated from a biological process, we seek to infer the true rates ***θ***^∗^ = (*λ*^∗^, *δ*^∗^, *µ*^∗^) governing the dynamics of division, death and mutation. Our goal is to maximize *P* (*A* ***θ***) with respect to the rates, thereby obtaining their maximum likelihood estimate (MLE) 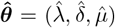. Since we lack an analytic expression for the likelihood of *A*, we use an expectation maximization (EM) algorithm, in which the backbone tree *B*, encoded by the node depths *s* and mutation times *m, t*, constitute our hidden variables, to iteratively find 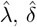 and 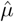.

To motivate our choice to use EM recall that for an allelic tree *A*, there is an infinite number of backbones *B* ∈ *f* ^−1^(*A*) satisfying the equation *A* = *f* (*B*). The probability of *A* given the rates can be formally written as

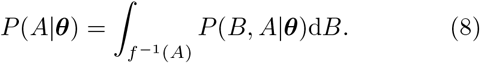

Note that since *A* is entirely determined by *B* through *A* = *f* (*B*), we have that *P* (*B, A* |***θ***) = *P* (*B* |***θ***). The backbone *B* in (8) contains the times at which mutations and divisions occur, i.e. hidden variables which are in theory integrated over to obtain the left hand side of the equation. In practice however, (8) cannot be computed analytically, and involves too many dimensions in the integral to be realistically computed numerically. We therefore use an expectation maximization (EM), a two-step iterative algorithm, to approximate the MLE.

In the first step, referred to as the “expectation” step, we calculate the expected value of the log likelihood of ***θ*** taken with respect to the current parameter estimates ***θ***_*i*_:

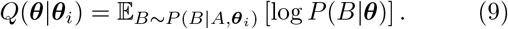

In the maximization step, *Q*(***θ*** | ***θ***_*i*_) is maximized with respect to ***θ***, yielding the parameter estimates for the next iteration

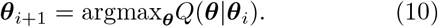

The log-likelihood term inside the expectation can be computed for each backbone tree using (1). However, the expectation value requires integrating over all backbone trees *B*, which is not practical. Instead, we evaluate that expectation value using Markov chain Monte Carlo (MCMC), which only requires to be able to compute *P* (*B*|*A*, ***θ***_*i*_) ∝ *P* (*B*|***θ***_*i*_) up to a constant of *B*. Specifically, we generate a sequence of samples using the Metropolis Hastings algorithm. Given a backbone *B*, a new backbone *B*^′^ ∈ *f* ^−1^(*A*) is proposed by sampling the depth or time of a random node or mutation from a uniform distribution with the condition that *f* (*B*^′^) = *A. B*^′^ is then accepted with probability

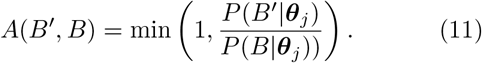

Starting from a random initializing backbone *B*_0_, we run the Metropolis Hastings algorithm for a burn-in period of *N* = 50 × (2*n* + *ℓ* − 1), where 2*n* + *ℓ* − 1 is the total number of variables *s, t, m* in *B*. After this period, we sample *k* backbones (*B*_1_, …, *B*_*k*_), and we approximate (9) by

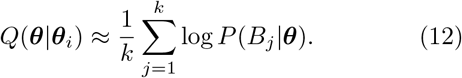

A total of *N*_MC_ expectation maximization steps are performed, yielding a series of estimates 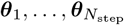. Since at each expectation step we only sample a finite number *k* of backbones, we do not expect the algorithm to converge exactly to the maximum likelihood estimate ***θ***_MLE_. Instead the estimates ***θ***_*i*_ are expected to fluctuate about ***θ***_MLE_ after a relaxation period *N*_*f*_, which empirical evidence suggests is short. We report our estimate of the parameters as an average over ***θ***_*i*_ following that relaxation:

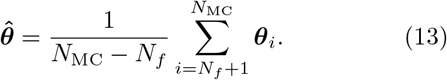

We can generalize the inference procedure to take more than one allelic tree as input. Consider a set of allelic trees (*A*_1_, *A*_2_, …, *A*_*N*_) all generated by processes with the same true rates ***θ***^∗^. The likelihood of the allelic trees then becomes

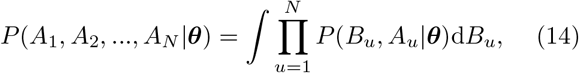

and the expected value of the log likelihood of ***θ*** must be taken with respect to all backbones

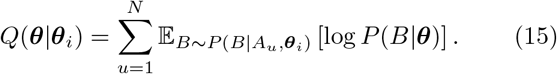

We create different independent realizations of our sampling scheme for each allelic tree, obtaining for each *A*_*u*_ a set {*B*_*u*1_, *B*_*u*2_, …, *B*_*uk*_} sampled from *P* (*B*| *A*_*u*_, ***θ***_*i*_). The approximation to equation (15) is given by

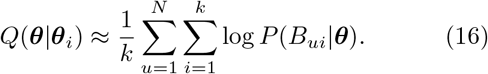

The maximization step does not change.

### D. Validation of inference procedure

We test our inference method on synthetic allelic trees generated with true parameters (*λ*^∗^, *δ*^∗^, *µ*^∗^), and with comprehensive sampling (*C* = *C*_*f*_). Practically, we generate extant cell lineage trees *C* using the coalescent point process (CPP) (Fig S1) [32], and apply the function *a* : *τ* → *A* to each of them to create a synthetic dataset of allelic trees *A*. The generation process along with a brief introduction to the CPP is detailed in Appendix 1.

We first test the inference for a single parameter given the true values of the other two and the total time *T*, beginning with the birth rate *λ*. In Fig. 4a, we show the distribution of birth rates inferred from a set of *N* allelic trees, where *N* is varied from 1 to 10, with *λ*^∗^ = 2.5, *δ*^∗^ = 1, *µ*^∗^ = 1.8, *T* = 3, and *k* = 1. Although the method theoretically is valid for large *k*, when the learning rate is slow, the ensemble average over *k* is emulated by an average over many learning steps around convergence. We checked that increasing *k* up to 10 did not affect the results of the inference (Fig. S4).

**FIG 4:**
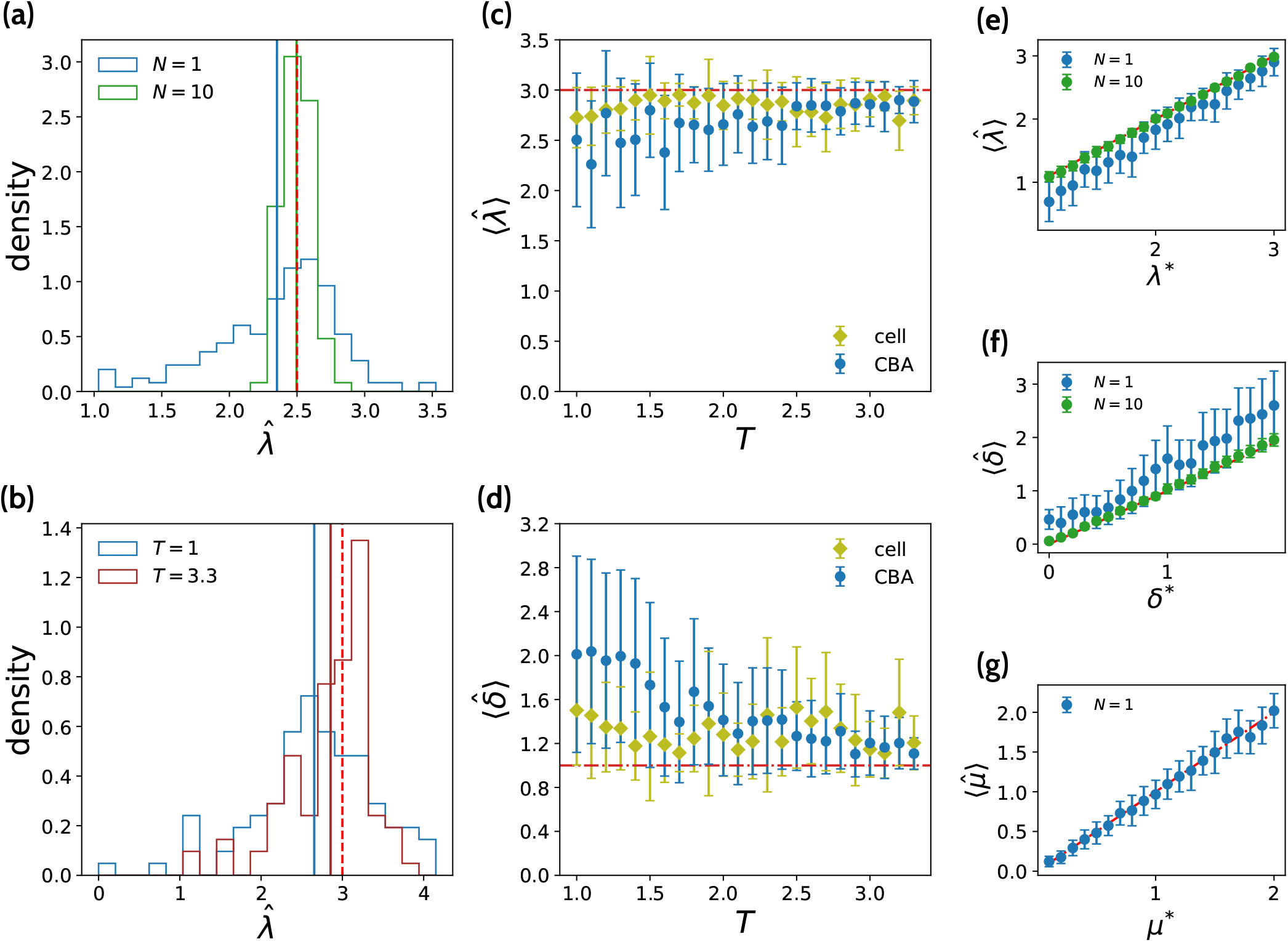
Validation of the inference method on synthetic data. Trees are generated with true parameters *λ*^∗^, *δ*^∗^, *µ*^∗^. The algorithm is used to infer one of the parameters, given the true values of the other two. Each numerical experiment is repeated 100 times to collect statistics. (a) Distribution of 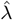 inferred from *N* = 1 (blue) or 10 (green) trees for *λ*^∗^ = 2.5 (dashed red line), *δ*^∗^ = 1, *µ*^∗^ = 1 and *T* = 3. Horizontal lines show 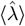. Estimates are less biased and more precise for larger *N*. (b) Same as (a), but with *λ*^∗^ = 3, *δ*^∗^ = 1, *µ*^∗^ = 1, *N* = 1 and two values of *T* : *T* = 1 (blue), and *T* = 3.3 (purple). Longer times lead to larger trees, which allow for better inference. (c,d) Average inferred (c) 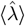 and (d) 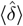and their standard deviations (represented by error bars) vs. *T*, using either inference on the allelic tree *A* (in blue), or the extant cell tree *C* (yellow). In (c) *δ*^∗^ = 1 and *µ*^∗^ = 1 while in (d) *λ*^∗^ = 3 and *µ*^∗^ = 1. The bias goes down as the tree size increases with *T*. This bias is still present when using the full information of the extant cell tree, although it is smaller. (e) Average inferred birth rate 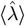 vs its true value *λ*^∗^ while keeping *δ*^∗^ = 1, *µ*^∗^ = 1 and *T* = 3 constant, for *N* = 1, 10. The bias goes down as *λ* and thus the size of the tree increases. It disappears for *N* = 10. (f-g) Same for the death and mutation rates, keeping *λ*^∗^ = 2, *T* = 3 and *δ*^∗^ = 1 while varying the mutation rate, and *T* = 3, *µ*^∗^ = 1 while varying the death rate. The bias is lowest when *δ* is small, and the trees the largest. There is no apparent bias in 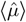even at *N* = 1. Error bars are given by the standard deviation of multiple inferences.

On average, when *N* = 1 (blue histogram), the inferred birth rate is very noisy and underestimated, i.e. 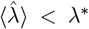. The large variance is a result of the large variability in the number of leaves of sampled extant cell trees, combined with fluctuations in both the number of observed branches and their lengths due to the random spread of mutations on *C*. Both bias and variance are due to finite sized effects, since maximum likelihood estimates are only guaranteed to converge asymptotically to the true parameter values in the limit of infinite data) *N*→ ∞. Indeed, 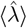gets much closer to the true value *λ*^∗^ when *N* is increased to 10 (green).

In our context, the large data limit may be directly achieved in two ways: by pooling more trees together (large *N*), or effectively by increasing the size of each tree, and with it the number of informative birth and mutation events that the MLE can exploit. The number of leaves *n*_*C*_ in the tree *C* of extant cells is on average:

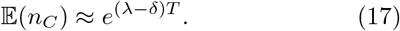

For a given mutation rate, the number of types in an allelic tree is heavily influenced by the number of leaves *n*_*C*_ in the extant cell tree it came from. In turn, the number of types is linked to how much information the allelic tree *A* carries about the parameters. Trees with more leaves contain more information and thus lead to better estimates for the rates. It is reasonable to consider the number of leaves in *C* as a proxy for the signal in *A*.

According to (17), larger trees may be obtained by either increasing the difference *λ* − *δ*, or by increasing *T*. Fig. 4b shows the distribution of inferred 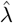 for *λ*^∗^ = 3, *δ*^∗^ = *µ*^∗^ = 1, and *T* = 1 (blue) or *T* = 3.3 (purple) with *N* = 1. As expected, much better and less biased inference is achieved on larger trees (*T* = 3.3) than on smaller ones (*T* = 1). We then plotted the average inferred 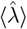, as well as its standard deviation, as a function of *T* (Fig. 4c). While *λ*^∗^ is always underestimated, the bias and standard deviation decrease as a function of *T*. Conversely, 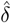 get overestimated, and that bias as well as the uncertainty also shrink with *T* and the tree size (Fig. 4d).

To investigate the origin of these biases in 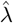 and 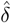, we asked whether inference with full information from the extant cell tree would also suffer from a similar bias. To do so, we used the original cell trees *C* from which the synthetic allelic trees were generated (and which are known since we’re dealing with synthetic data), and inferred the rates *λ* and *δ* with a maximum likelihood estimator, using analytic formulas for the probability *P* (*C*) of the realized cell tree from the CPP [32]. Details of the used formulas are given in Appendix 1. Results for the average inferred parameters and their standard deviations are given by the yellow curves in Fig. 4c and d. They show that the bias still exists, albeit less so than for inference on the allelic tree. It also decreases as the tree size grows by increasing *T*. The smaller bias and variance are consistent with the fact that cell trees contain more information than allelic trees, including mutation and division times as well as abundances of cells of the same type. However, the fact that the bias is still present points to a more fundamental origin.

In fact, the overestimation of *δ*, and related underestimation of *λ*, may be understood intuitively by considering a simpler case, where the observation is simply the number of leaves *n*_*C*_ in *C*. In that case, the MLE estimator of *λ* would be 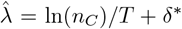, and the MLE estimator of *δ* would be 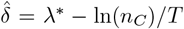 (see SI Appendix **??** for a derivation). Since ln ⟩ (*n*_*C*_)) ≤ ln *n*_*C*_) ⟩ = (*λ*^∗^ − *δ*^∗^)*T* by Jensen’s inequality, this implies 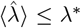 and 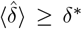, consistent with our observations. While the inference on the full *C* or on the backbone tree *B* contain different sorts of information than just *n*_*C*_, such an effect may affect our MLE estimator. Inspired by the previous argument, we shown in Fig. S5 that when estimating 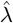or 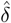with CBA, calculating 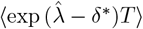 or 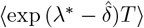 provides an unbiased estimate of the average number of leaves (*n*_*c*_).

We then study how the inference performs as a function of the parameters. We vary *λ*^∗^ across a given range, keeping *δ*^∗^ = 1 and *µ*^∗^ = 1 constant and known, and estimate the average MLE inferred 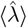 for a large number of groups of *N* trees as a function of *λ*^∗^, for *N* = 1 and *N* = 10 and *T* = 3 (Fig. 4e). For *N* = 1, *λ*^∗^ is consistently underestimated, with smaller *λ*^∗^ leading to both larger biases in 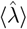 and higher uncertainty. This is expected, since for constant *δ*^∗^ and *µ*^∗^ a higher *λ*^∗^ leads to more larger and more informative trees. Increasing the number of trees to *N* = 10 for each inference is sufficient to remove the bias in (*λ*) (Fig. 4e).

To better understand how the amount of data impacts the accuracy of the estimates, it is interesting to compare the cases of a single large tree (*λ*^∗^ = 3.0, *N* = 1) to many small trees (*λ*^∗^ = 1.1, *N* = 10), all with *µ* = *δ* = 1 and *T* = 3. We have *n*_*C*_(*λ* = 1.1) ≈ *e*^0.3^ ≈ 1.35 and *n*_*C*_(*λ* = 3) ≈ *e*^6^ ≈ 403. We would expect the signal in a single tree with *λ* = 3 to be much larger than 10 times the signal in a tree with *λ* = 1.1. However, Fig. 4b shows that we obtain a better estimate of the birth rate when *N* = 10 and *λ*^∗^ = 1.1. This indicates that the number of allelic trees used for the inference is much more important than their sizes.

We also varied the other parameters, by applying maximum likelihood to infer a range of *δ* and *µ*. We vary *δ*^∗^ ∈ [0, 1.9], keeping *λ*^∗^ = 2 and *µ*^∗^ = 1 fixed, and *T* = 3. Fig. 4f shows the relationship between 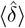 and *δ*^∗^. Inferring the death rate of reconstructed phylogenetic trees is generally considered a harder task than inferring the birth rate, since dead cells are not represented in the data. Comparing the results for *N* = 1 in Figs. 4b and 4c, it is clear that 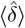 is more biased and more noisy than 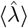. Consistent with previous observations, 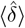 is consistently overestimated, although that bias disappears for *N* = 10. Finally, in Fig. 4g we show the average inferred mutation rate 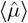 at fixed *λ*^∗^ = 2 and *δ*^∗^ = 1 for *N* = 1. In that case, a single allelic tree is sufficient to obtain an unbiased estimator for the mutation rate. We also verified that inference of other parameters, such as *λ*, became for more precise and less biased as the mutation rate is increased (Fig. 5). Indeed, we expect the backbone tree to provide a good approximation of the extant cell tree when the mutation rate is large.

**FIG 5:**
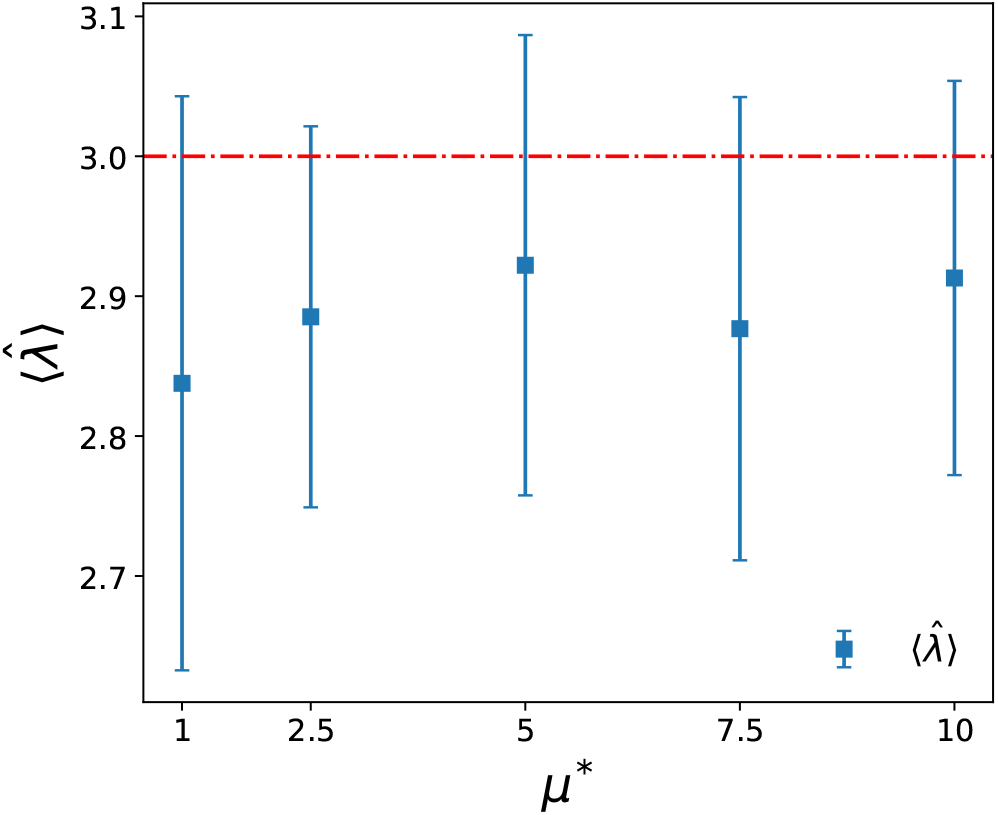
Dependence of birth rate inference on the mutation rate. Average 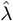 inferred as a function of the mutation rate for *λ*^∗^ = 3 (dotted red line), *δ*^∗^ = 1, *T* = 3 and *N* = 1. Larger mutation rates imply larger allelic trees *A* allowing for more accurate inference. As *µ*^∗^ increases we note an increase in 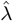 and a decrease in the error. Numerical experiments are repeated 200 times to gather statistics. Error bars are given by the standard deviation of multiple inferences.

In most situations only a small fraction *ρ* of cells are sampled and used to construct an allelic tree. In that case, our inference formulas (1)-(7) are still valid, but with an additional dependency of the *p*(*t*) and *q*(*t*) function on *ρ* (see Appendix 1 and 2). Performing the inference on subsampled trees with *ρ* = 10% still works, although accuracy is slightly degraded, as expected from the loss of information caused by subsampling (Fig. 6).

**FIG 6:**
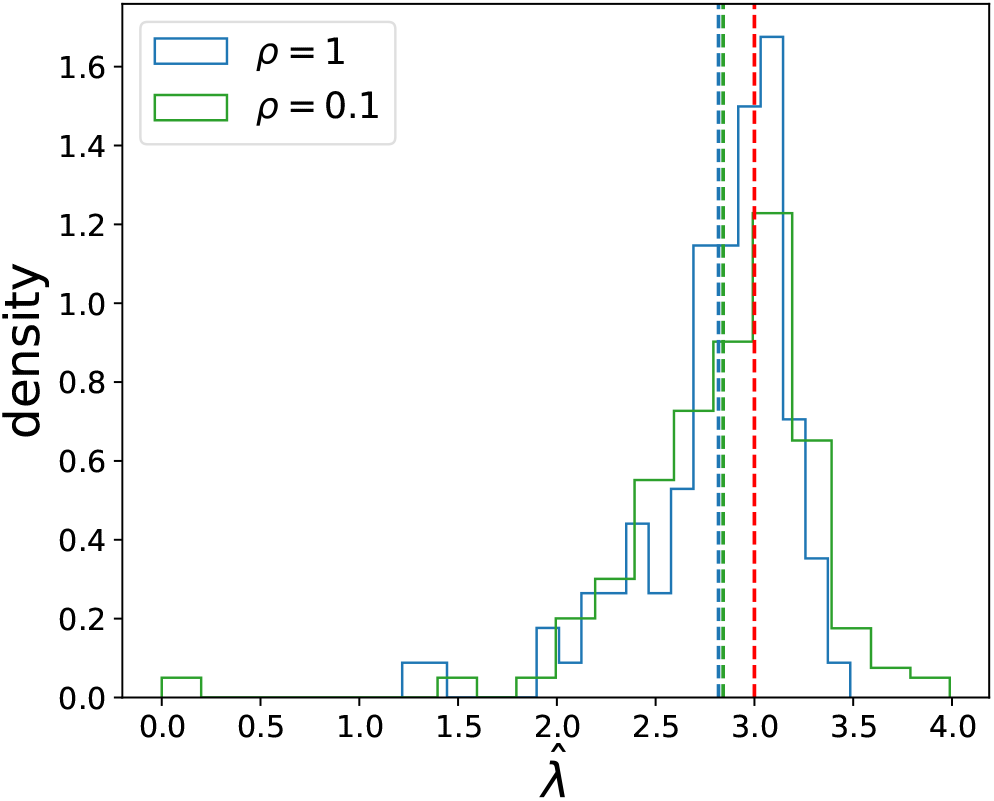
Inference with sampling. Distribution of 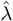 inferred from *ρ* = 1 (blue) and *ρ* = 0.1 (green) for trees generated with *λ*^∗^ = 3 (dotted red line) and *N* = 1, given *δ*^∗^ = 1, *µ*^∗^ = 1 and *T* = 3. No significant change in the average birth rate 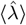 (dotted lines) is observed when under sampling. We note that the distribution is wider when *ρ* = 0.1, indicating noisier inference. The distributions are the results of 200 repetitions of the numerical experiments.

Next, we consider the task of simultaneously inferring *λ* and *δ*, given *µ*^∗^ and *N* = 1 and *T* = 3. To get a sense of the difficulty of the task, we first study inference on synthetic backbone trees. Our method is ultimately designed to make inference on allelic trees, but since backbone trees contain more information, we reason that issues observed in the inference on backbone trees will be inherited by allelic trees. In Figure 7a we show a heat maps of the likelihood *P* (*B* | *λ, δ, µ*^∗^) of a backbone tree *B* as a function of *λ* and *δ*, where *B* was sampled from *P* (*B*| *λ*^∗^, *δ*^∗^, *µ*^∗^) with *λ*^∗^ = 3, *δ*^∗^ = 1, *µ*^∗^ = 1 and *T* = 3. The likelihood has a well defined maximum, but is flat in the difference *r* = *λ* − *δ*. This implies that the effective growth rate *r* is easier to infer than the individual rates *λ* and *δ*.

**FIG 7:**
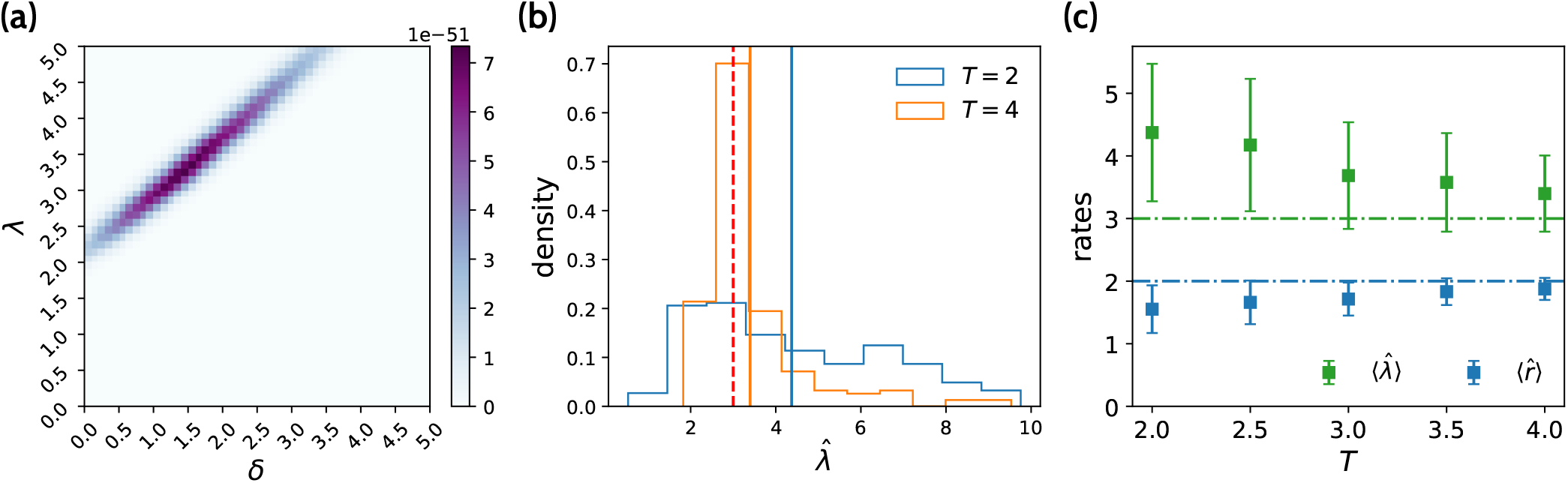
Two-dimensional inference. (a) Heatmap representing the normalized likelihood *P* (*C* ***θ***)*/ ∫ d****θ****P* (*B* ***θ***) as a function of two unknown parameters *λ* and *δ* for *N* = 1 with known *µ*^∗^ = 1 and *T* = 3, for a backbone tree *B* drawn with *λ*^∗^ = 3, *δ*^∗^ = 1. The likelihood landscape shows that while the difference *r* = *λ* − *δ* is well represented by the tree, the value of each *λ* and *δ* are very uncertain. Note that this inference on *B* is different from the inference on *A* we are interested in, but it allows for an explicit computation of the likelihood which our method on *A* does not permit, giving us insight by proxy. (b) Distribution of inferred 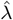 in the two-dimensional EM search for 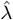 and 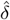, with the same parameters as above, but for *T* = 2 (blue) and *T* = 4 (orange). Longer times rates allow for more accurate inference. (c) Mean 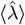 and 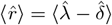 as a function of age *T*, for the same parameters as above. Note that the effective growth rate 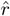 is underestimated, consistent with previous observations on the single-parameter inference. In contrast, 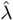 is now overestimated. For each *T* we perform 200 independent simulations to gather statistics. Error bars are given by the standard deviation of multiple inferences.

We then applied the EM algorithm to infer both *λ* and *δ* simultaneously from allelic trees. As *T* is increased, estimates for *λ* and *µ* improve both in bias and in variance. This is evidenced by Fig. 7b, where the distribution 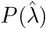 is shown for *T* = 2 and *T* = 4, as well as by Fig. 7c, in which we plot the dependence of 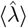 and 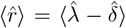 on *T*. Curiously, the birth rate *λ* is consistently over-estimated, in contrast to the single-parameter inference case. However, the effective growth rate *r* = *λ − δ* is still underestimated, consistent with the convexity argument above.

### E. Comparison to other methods

We are not aware of other methods specifically designed to infer rates from a single tree with few mutations. However, existing software with broad applicability can be utilized to perform that task. We compare our results to those obtained using the Bayesian evolutionary analysis by sampling trees (BEAST) [20] software for the task of simultaneously inferring the birth and death rates *λ* and *δ*, given the mutation rate *µ*^∗^. To do so, we first need to reformulate our problem so it can be handled by BEAST. In contrast to our inference scheme which begins from the allelic tree *A*, BEAST takes as input a multiple sequence alignment (MSA) ℳ. It then samples from the posterior *P* (***θ***, *A*^′^ |ℳ) ∝ *P* (ℳ|***θ***, *A*^′^)*P* (***θ***, *A*^′^), i.e. the joint distribution of parameters ***θ*** and trees *A*^′^givenℳ, using a Metropolis-Hastings algorithm. The sampled trees *A*^′^ differ from our allelic trees in two ways, as illustrated in Fig. S3. *A*^′^ is time-resolved, meaning that multifurcations are timed and broken up into bifurcations. Also, the root of *A* (green node in Fig. S3) is not contained in *A*^′^, and *A*^′^ starts with the most recent common ancestor.

To generate the input data ℳ usable by BEAST, we first generate an allelic tree *A*, and create a set of synthetic sequences corresponding to the leaves of *A*, and consistent with its topology (see SI 4). In addition, we specify priors on all the different subsets of sequences which have a common ancestor in *A* to ensure that all the trees sampled during the inference performed by BEAST are constrained to have the same topology as *A*. The mutation rate *µ*^∗^ is fixed and hardcoded into BEAST, and the prior over the birth and death rates are chosen so that *r* = *λ* − *δ* is uniformly distributed between 0 and 1000, and *δ/λ* uniformly distributed between 0 and 1. By feeding ℳ and subsets of leaves with common ancestors to BEAST, we infer the estimates 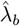 and 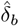 as the mean values from the posterior 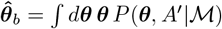. These estimates are then compared to the 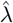 and 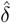 inferred using CBA on *A*.

Fig. 8 shows a comparison of our MLE estimator with the estimates of BEAST. We observe that for small mutation rates, our approach is consistenly unbiased, while BEAST systematically underestimates the birth rate even at large mutation rates, and only converges towards the true value of the parameters as *µ* becomes large. Since BEAST was designed for genetic data where types differ by many mutations, it is expected to perform best in the limit of large numbers of mutations, where the branch lengths are a good approximation of the times between speciation events. At low mutation rates however, where ancestral types are sometimes represented in the data and branch lengths correlate only weakly with time, our methods performs better.

**FIG 8:**
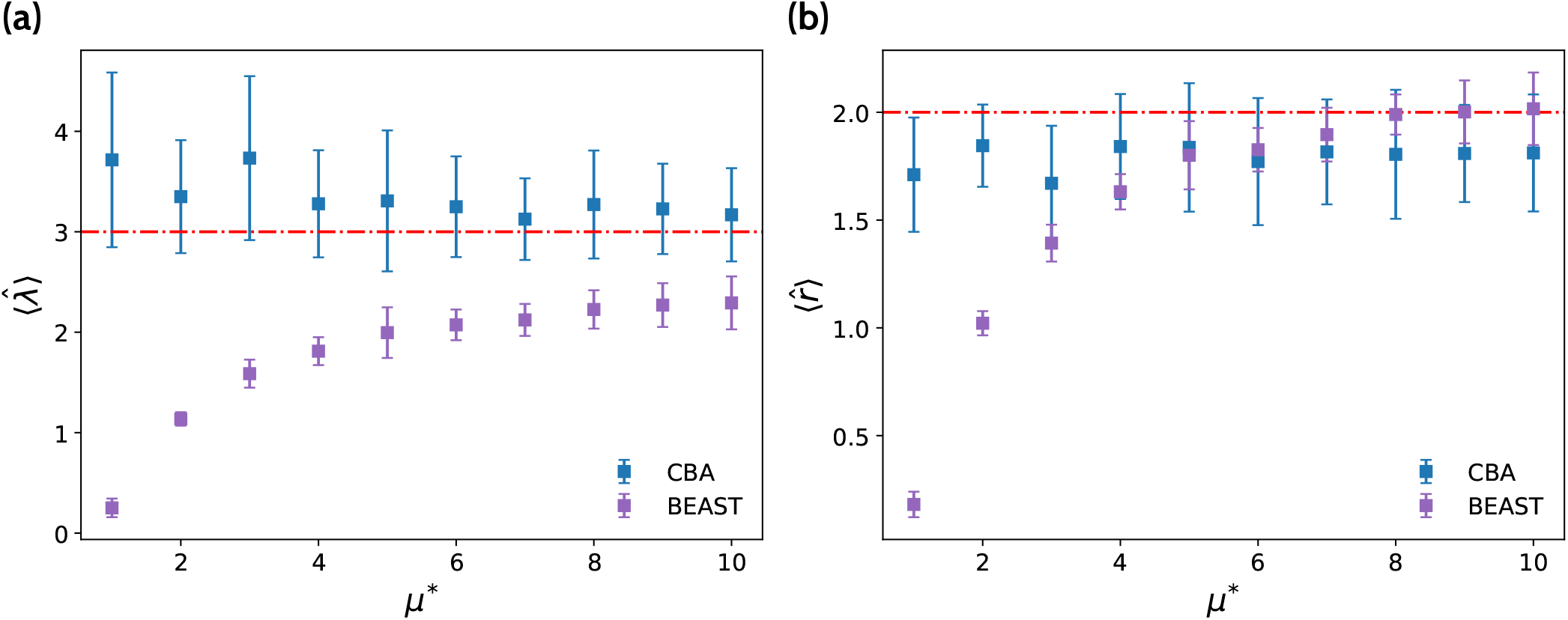
Comparison with BEAST [20]. Average inferred (a) birth rate 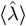 and (b) effective growth rate 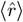 as a function of the mutation rate *µ*^∗^, for both our method and the posterior average in BEAST, in the two dimensional inference of *λ*^∗^ = 2.8 and *δ*^∗^ = 1 with known *µ*^∗^, from a single allelic tree *A* (*N* = 1). In order to keep the size of the allelic trees comparable across mutational regimes (number of types *≈* 100), the stem age of the tree *T* is decreased for increasing *µ*^∗^. Our method performs best compared to BEAST at low mutation rates, where branch lengths are a poor approximation of time. At large mutation rates, both methods perform well. In both (a) and (b) error bars are given by the standard deviation of 100 independent inferences.

## III. DISCUSSION

We introduced a novel method designed to infer evolutionary rates of a growing population of cells sampled at a single time point from an allelic tree *A* describing mutational histories. We derived the likelihood of the backbone tree *B*, which enabled us to come up with an efficient inference scheme based on expectation maximization to find the maximum likelihood estimates of the rates. We limited our analysis to rates ***θ*** = (*λ, δ, µ*) that were homogenous in time, which allowed for the separate inference of *λ* and *δ*. All the formalism and derivations introduced are easily generalizable to rates with an explicit time dependence ***θ***(*t*) = (*λ*(*t*), *δ*(*t*), *µ*(*t*)) (see [32] for a generalization of equations 18 - 19). However, doing so would require parametrizing these dependences with additional parameters, requiring more data to support the inference.

We derived the method in a general framework, since applying it to concrete systems often requires checking the validity of the assumptions. Firstly, we assumed that mutations accumulate continuously over time, instead of upon birth. Both situations occur in cells. Conversely to what we assumed somatic mutations occur during DNA replication, however they can also occur continuously throughout the cells cycle due to DNA repair errors which lead to mismatches that are incorrectly resolved [37]. On the discrete side, some single-based substitution (SBS) signatures are known to increase with cellular turnover (e.g. SBS1) [38], indicating that errors in DNA replication occurring at discrete times are responsible. Yet other signatures are thought to be mainly caused by errors in DNA repair, owing to their quasi-continuous clock-like accumulation in post mitotic cells [37]. In the example of affinity maturation, B-cells somatically hypermutate when the Activation-Induced cytidine Deaminase (AID) deaminates a cytidine to a uracil, creating mismatches which can be incorrectly repaired by the error prone DNA repair mechanism [39]. These processes usually are assumed to accumulate continuously, although recent work shows that it can depend on the cell cycle [40]. Others have made both continuous and discrete time assumptions for haematopoietic stem cells. The cloneRate inference method [22] infers the net growth rates of cell colonies derived from single haematopoietic stem cells [13, 14, 41, 42] under the same assumption that mutations accumulate through time at a constant rate, while another recent method [25] analyzes two haematopoeisis phylogenies from neonate donors [14] under the assumption that mutations occur upon division with a Poisson distribution.

In contrast to BEAST [20], we do not discuss the task of inferring *A* from a multiple sequence alignment (MSA) — an active field of research [4, 30, 43, 44] — as our inference scheme starts from the allelic tree *A*. We thus assume that data is in a regime where the inference of *A* is accurate with no degeneracy. We expect this assumption to be true when the total number of mutations is low compared to the length of sequences in the MSA (mutation frequency). For example, the mean nucleotide mutation frequency in B-cell receptor V genes is known to be approximately 5% both in healthy and chronically HIV-1 or HCV-infected individuals [45, 46], which should garantee reliable allelic tree inference. In addition, a likelihood-based method incorporating genotype abundances, obtained from single-cell data, was shown to accurately lift the degeneracy in B-cell phylogenies inferred using maximum parsimony [4]. In haematopoetic stem cells whole genome sequencing shows that they accumulate on average 17 somatic mutations per year [41]. An 80 year old individual would thus have an approximate mean nucleotide frequency of 0.00005%, assuming ∼3 billion base pairs in the genome, making inference of *A* unambiguous.

Most evolutionary processes are subject to selection and a lot of work has gone into identifying signatures of selection. Cancers are caused by the accumulation of driver mutations which confer selective advantages to cells in which they appear, while viral-immune coevolution places selective pressure on both viruses and the immune system [47, 48]. Methods based on the ratio of synonymous to non synonymous mutations (d*N/*d*S*) can efficiently identify the average number of driver mutations in different tumors [19]. For the seasonal A′H3N2 viruses, assuming evolution proceeds through the continual accumulation of weakly deleterious or beneficial mutations, the branching patterns of time resolved phylogenies can be used to infer the fitness of the sampled strains [24]. Many viral and macro-evolutionary phylogenies benefit from branches containing a large number of mutations, allowing for fitness dynamics to be approximated by selection biased diffusion [24]. In the case of B cell immunoglobulin (Ig) receptors, due to the low average number of mutations per sequence [49], this approximation cannot be made. Recent efforts in quantifying selection have focused on comparing the frequency of non-synonymous mutations to their expected frequency assuming neutrality [7, 8], with the latter being difficult to calculate. Inferring signatures of selection from allelic trees has been done for B cells sampled from individuals subjected to the trivalent influenza vaccine [10] or with chronic HIV-1 infections [9] by focusing on statistics such as tree branching asymmetry, terminal branch length distributions and the site frequency spectrum. Multitype birth death processes (MTBD) model selection by allowing for birth and death rates to change discretely along phylogenies [50]. They have been used for Bayesian inference in macroevolution [50], whereby time-resolved phylogenies are sampled along with rates using molecular clock models. We are not aware of any existing methods which infer evolutionary rates using MTBD models for allelic trees when the molecular clock hypothesis breaks down, e.g. for B cell phylogenies. The generalization of our model past neutrality suffers from our use of coalescent point processes to derive the likelihood of the backbone, specifically through the terms *p*(*t*) and *q*(*t*) (equations 28 and 30). The chief condition for reconstructed trees of birth-death models with rates *λ*(*t*) and *δ*(*t*) to be CPPs is that while the rates may depend on absolute time *t*, they cannot be subtree dependent. Clearly, this assumption does not hold when selection is present, since the descendants of a cell with a higher birth rate than its ancestor will carry this change in fitness along the subtree. A possible extension for future work would be to use dynamic programing [24] or to use approximations to compute the likelihood of *B* accurately.

Besides the issue of selection, a direct application to B cells would require choosing appropriate time dependence for the rates ***θ***(*t*) = (*λ*(*t*), *δ*(*t*), *µ*(*t*)). B cell affinity maturation takes place in germinal centers and is spatially heterogeneous [39]. Germinal centers are divided into dark zones (DZ), where B cells somatically hypermutate and proliferate, and light zones (LZ), where B cells are selected for their affinity to an antigen [39]. Cells migrate independently between the two zones in a process known as cyclic reentry, with migration rates in mice depending on whether cells are transitioning from the LZ to the DZ or vice versa [51]. Additionally, the low percentage of cells which reenter the dark zone indicates the presence of strong bottlenecks where most cells die or go away [51]. To model cyclic reentry, *λ*(*t*) and *µ*(*t*) could be defined as piecewise functions alternating intervals in which birth is high and where death is high. Increasingly complex choices for ***θ***(*t*) could lead to non identifiable statistical models [52], highlighting the importance of simple parametrizations.

In our numerical experiments we found a consistent bias in the estimation of *λ* and *δ*, which disappears asymptotically in the large-data limit. In the single parameter inference from a few small trees, *λ* was over-estimated and *δ* was underestimated. When inferring *λ* and *δ* simultaneously, we found that both parameters were overestimated, leading to an underestimation of their difference *r*. To circumvent this bias, we could consider extending CBA to Bayesian inference. Through clever choices for the priors over the rates *P* (*λ*), *P* (*δ*) and *P* (*µ*), sampling from the posterior *P* (***θ***, *B*| *A*) could lead to unbiased estimators as well as get rid of the need for the expectation maximization algorithm. Finally, in many experiments the time *T* at which cellular expansion began is unknown. However, prior knowledge on *T* can easily be integrated into the inference.

## Acknowledgements

This work was supported by CNRS 80 Prime Grant (AMW, TM, AL), the Agence Nationale de la Recherche grant no ANR-19-CE45-0018 “RESP-REP” (AMW, TM, PP) and the CZI Theory Initiative.

---

**FIG S1:**
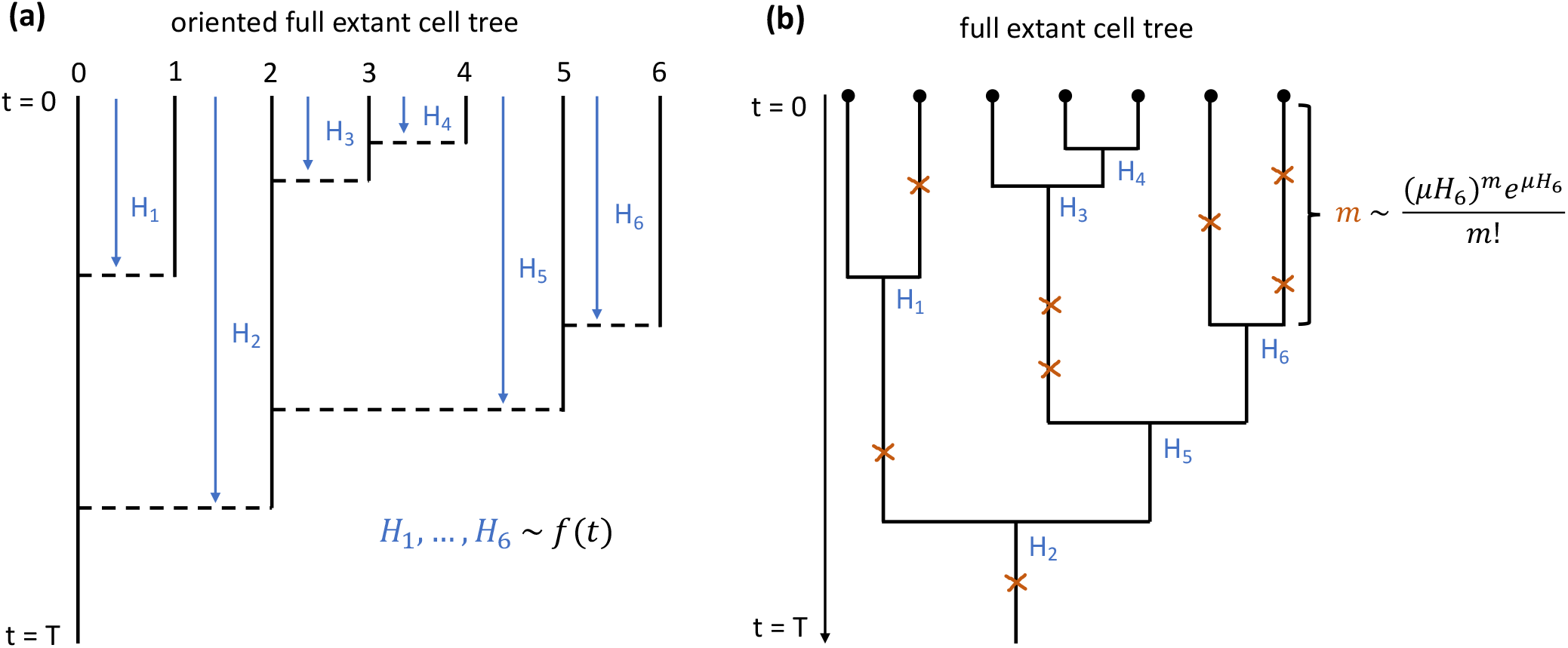
Generation of trees with CPP. (a) A coalescent point process with seven tips. Node depths are drawn from the coalescent density *f* (*t*) and placed in order in the plane until a node depth *H > T* is drawn. Horizontal dotted lines are drawn from the bottom of node depths until they reach a vertical line, thus completing an oriented reconstructed phylogenetic tree. (b) The non oriented reconstructed phylogenetic tree, obtained from (a) by ignoring the planar orientation but respecting the timing and order of coalescence events. Mutations are spread on the non oriented, reconstructed phylogeny by drawing, for each branch with length *ℓ* (e.g. *H*_6_), the number of mutations from a Poisson distribution with parameter *µℓ* (e.g. *µH*_6_) and placing them uniformly on the branch.

## Supplementary information

### 1. Generating allelic trees using coalescent point processes

Birth-death process are used to model a plethora of diverse phenomena, from queues and waiting lines in grocery stores to the evolution of biological species. In the context of macroevolution, birth-death models are generally assumed to have rates which are either constant, or which may depend on (1) absolute time, (2) the number of coexisting species, (3) a non-heritable trait and (4) a heritable trait. Models with rates depending on any or all of the 4 can be simulated forward-in-time using, for example, the Doob-Gillespie algorithm [53]. However, writing the likelihood of reconstructed phylogenies, i.e. phylogenetic trees of surviving species, generated by those birth-death processes is not feasible in all cases. In Lambert and Stadler (2013) [32] an analytic expression for the likelihood of such trees was found for birth-death models where the birth rate *λ* depends at most on absolute time *t* and where the death rate *δ* may depend on absolute time *t* and a non-heritable trait *x*. They showed that birth-death models of this kind are coalescent point processes (CPP), where a CPP of stem age *T* is an oriented full extant cell tree of depth 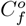 whose node depths *H* are independently and identically distributed according to the coalescent density

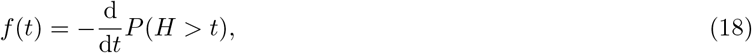

where *P* (*H > t*) is the tail distribution, i.e. the probability that a node depth *H* is greater than some time *t* (Fig. S1a). The orientation of 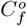 refers to the distinction made upon division between a parent (left branch) and its child (right branch). All oriented trees with *n* tips can give rise to 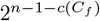 equally likely full extant cell trees *C*_*f*_, where *c*(*C*_*f*_) is the number of non symmetric nodes in the tree. The inverse tail distribution is given by

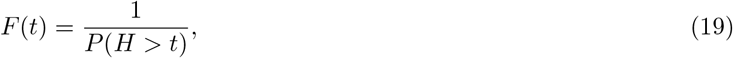

and can be used to rewrite Eq. 18 as

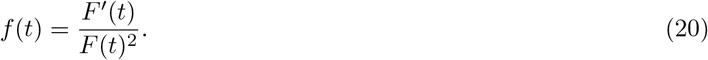

**FIG S2:**
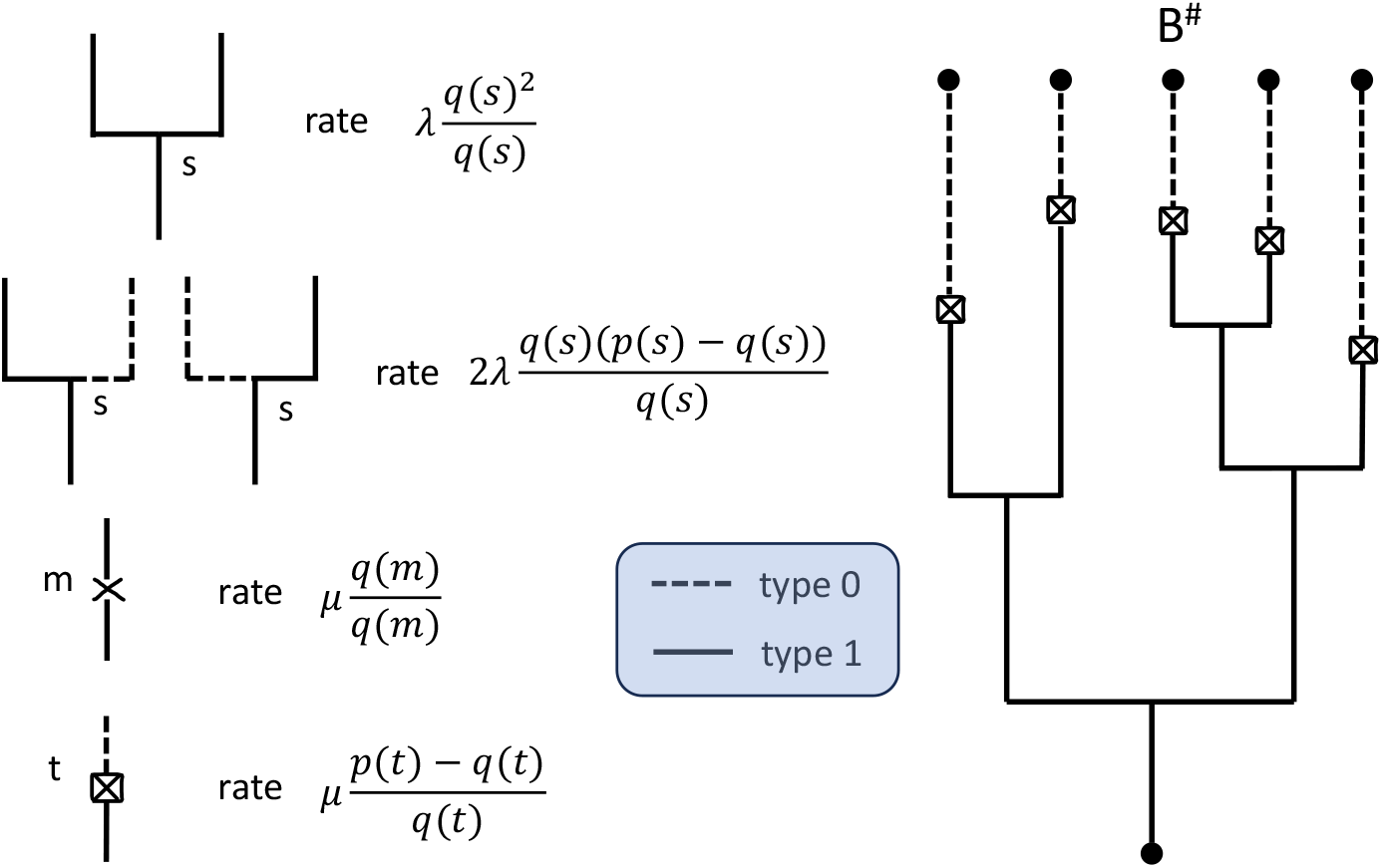
Splits and freezes. Type 1 (0) particles have (no) mutations in downstream subtrees. Particles of type 1 (solid lines) can split into two particles of type 1 or into one type 1 particle and one type 0 (dotted line) particle. Particles of type 1 can mutate and keep their type or mutate and freeze into a type 0 particle. The tree *B*^#^ is the backbone tree *B* with no internal mutations.

For constant rates *λ, δ, µ* and sampling parameter *ρ*, equation 19 becomes

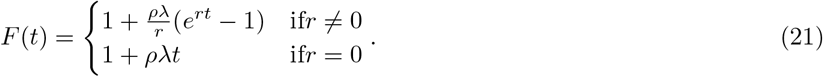

Consequently, we can write equation 18 as

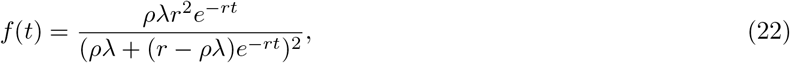

where *r* = *λ−δ*. The likelihood of a full extant cell tree *C*_*f*_ with *n* leaves and stem age *T* is given by the product over the coalescent densities of all the node depths, times the probability that the *n*th node depth is larger than *T*, i.e.

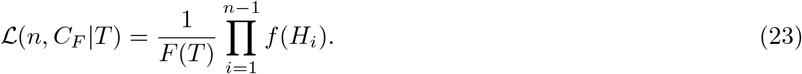

To generate an allelic tree, we first sample an oriented full extant cell tree 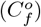 by following the CPP procedure [32], i.e. conditioned on a stem age *T*, sample node depths from *f* (*t*) until a value larger than *T* is drawn. We then pick a random non-oriented representation of 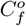 and spread neutral mutations on it through a Poisson point process (Fig. S1b). The number of mutations *m* on every branch is drawn from the Poisson distribution

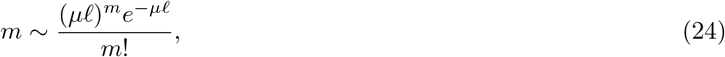

where *ℓ* is the branch length. The *m* mutations are then spread uniformly over the branch. The tree resulting from the process thus far is the full extant cell lineage tree *C*_*f*_. To collapse *C*_*f*_ into *A*, we apply the function *A* = *a*(*C*).

### 2. The likelihood of the backbone tree

We begin by recalling the definition of the full extant cell tree *C*_*f*_ as the cell lineage tree *D* with pruned extinct lineages. The cell lineage tree itself is generated by a birth-death process with neutral mutations, with birth rate *λ*, death rate *δ* and mutation rate *µ*. The backbone tree *B* is defined as the tree obtained from *C*_*f*_ where (1) subtrees downstream of terminal mutations are replaced by a branch (2) subtrees downstream of internal mutations are collapsed to only keep the part of the tree leading to further mutations and (3) boxing (circling) a mutation if there exists (does not exist) a path between the mutation and the present with no further mutations.

A tree stemming from one cell at depth *t* from the present has (1) extant descendants at the present with probability *p*(*t*) (2) extant descendants and mutations with extant descendants at the present with probability *q*(*t*) and (3) extant descendants integrally clonal to the cell with probability *p*(*t*) − *q*(*t*). To derive these probabilities, we make use of the coalescent point process formulation of full extant cell trees, introduced in Appendix 1. For *p*(*t*), we consider a portion of edge *e* connecting two nodes at depths *a* and *b*, with *b > a*. The probability *P*_*n*_ that along *e* no tree sprouts which survives to the present is given by

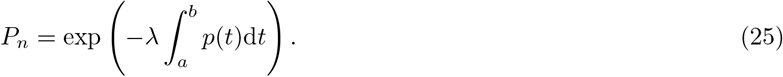

In the CPP picture, *P*_*n*_ is the probability that the next branch with depth larger than *a* has depth larger than *b*

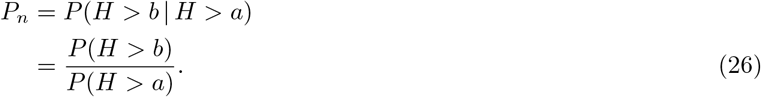

Equating the right hand side of Eq. 25 to the right hand side of Eq. 26 and recalling the definition of the inverse tail distribution we obtain

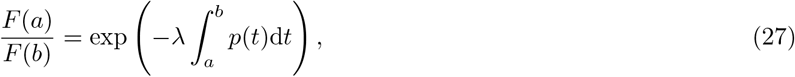

which, when differentiated with respect to *a*, gives

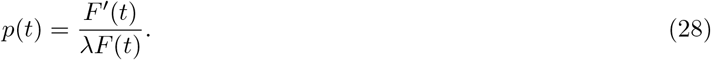

To derive *q*(*t*) we first solve for the probability that a full extant cell tree of depth *t* has no mutations, which can be written as

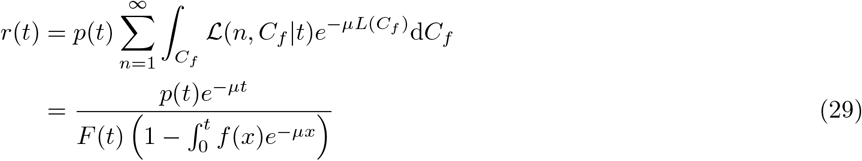

where *L*(*C*_*f*_) is the total length of *C*_*f*_ and L(*n, C*_*f*_ |*t*) is the likelihood of the oriented reconstructed cell tree. By recognizing that *r*(*t*) = *p*(*t*) − *q*(*t*) we obtain the expression for *q*(*t*)

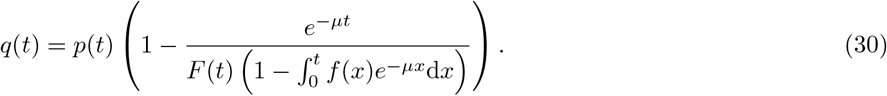

Returning to the derivation of the likelihood, we assume that the tree *C*_*f*_ has extant descendants and at least one mutation represented at the present, which happens with probability *q*(*T*). A particle is of type 1 (0) if it has nonclonal (integrally clonal), extant descendants at the present. Particles of type 1 may split into two particles of type 1 at rate

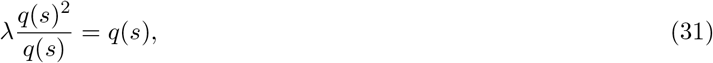

and into two particles of type 0 and type 1 at rate

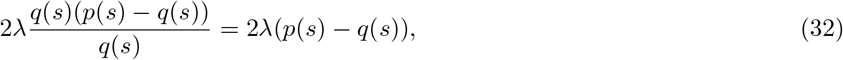

where we divide by *q*(*s*) because the particle is conditioned to be of type 1 before the split (Fig S2). The factor of two in equation 32 accounts for the different ways in which the branching can occur (type 0 can be on the left or right).

Particles of type 1 can mutate and keep its type at rate

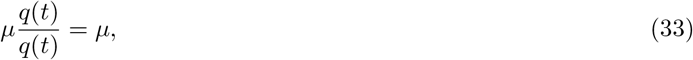

*q*(*t*) and will mutation and freeze into lineages of type 0 at rate

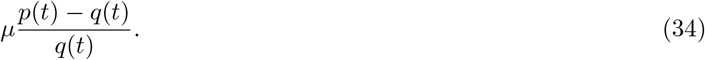

Particles of type 0 no longer divide or mutate.

Conditioned on survival of the birth death process and at least one represented genotype, the previous paragraph offers a prescription for simulating backbone trees. Start with one lineage of type 1 at time *T*. Lineages of type 1 split at rate *λq*(*t*) and freeze into lineages of type 0 at rate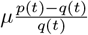. These splits and freezes occur until the present, so *t* = 0, generating an object we denote by *B*^#^. The likelihood of *B*^#^ is given by

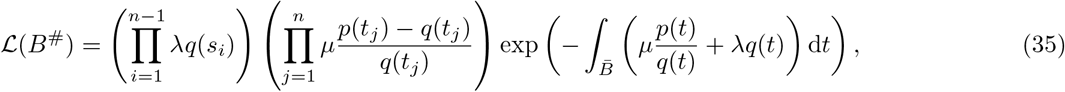

where {*s*_*i*_} are node depths, i.e. the times at which the branchings occur, {*s*_*i*_} are the depths of the leaf mutations, and 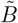 is the blue part of *B* ^#^, i.e. *B* ^#^ without the frozen edges. The exponential term in equation 35 accounts for the probability of no mutations or divisions occurring over the entirety of 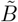.

Inner mutations occur on all lineages of type 1 at rate *µ* and lineages with clonal, extant descendants sprout at rate 2*λ*(*p*(*t*) − *q*(*t*)). Denoting by *L*_*i*_ the concatenation of intervals downstream of an internal mutation *i*, the mutation will be circled with probability

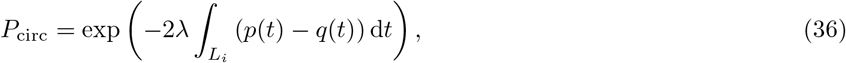

or boxed with probability

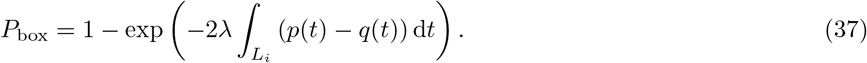

Now, conditioned on *B*^#^, internal mutations fall on the blue part of *B*^#^ as a Poisson point process with rate *µ* and each of these mutations *m*_*i*_ for *i* = 1, …, *r* are circled (boxed) with probability *P*_circ_ (*P*_box_). Therefore we can write the likelihood of *B* as

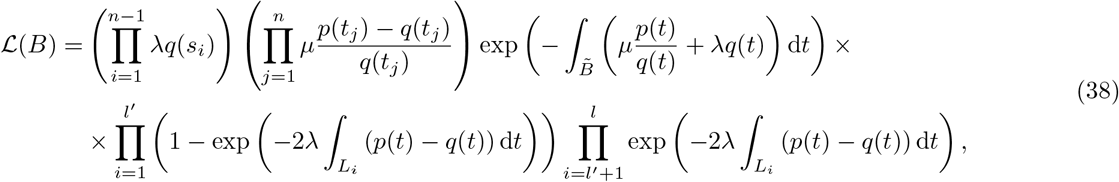

where *l*^′^ is the number of boxed mutations.

### 3. Maximum likelihood from the number of leaves in the tree of extant cells

The number of extant individuals in a birth-death process follows a geometric distribution of mean 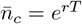 :

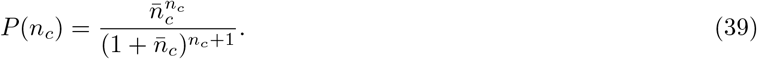

The MLE estimate is obtained by solving *∂* ln *P* (*n*_*c*_)*/∂r* = 0 for *r*, which gives:

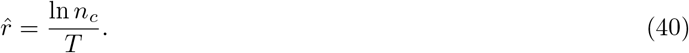

When doing inference on *λ* only with known *δ*, or vice-versa, give respectively:

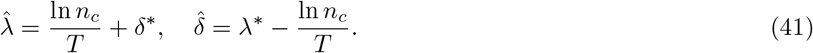

**FIG S3:**
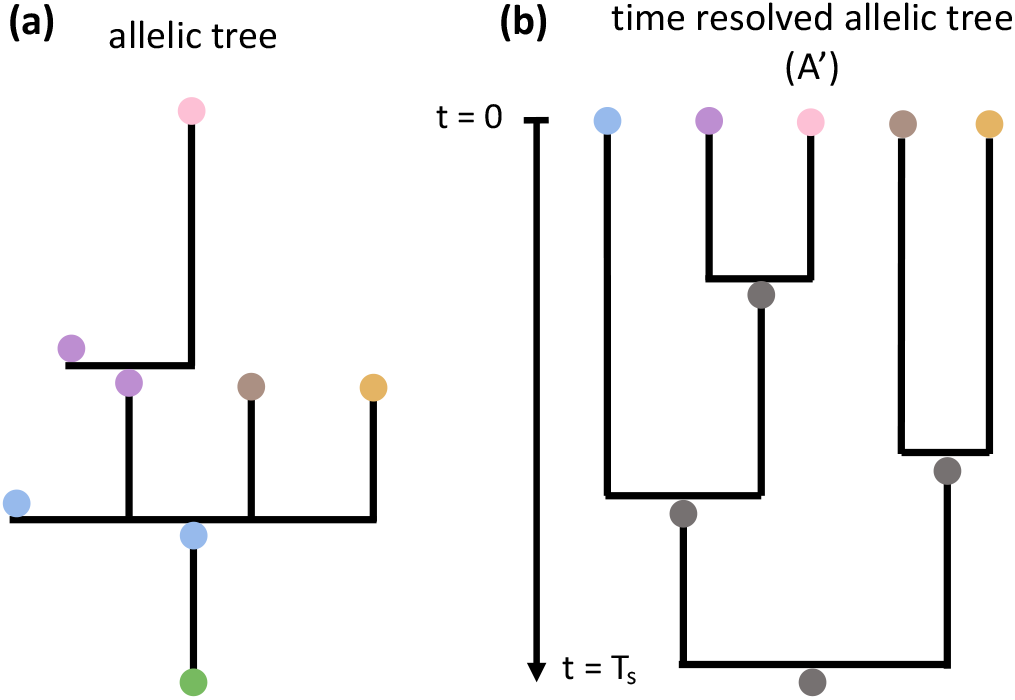
BEAST trees. (a) An allelic tree A (b) The time resolved allelic tree A’ corresponding to the allelic tree in (a) where the multifurcations are timed and resolved into bifurcations. A’ starts with the most recent common ancestor, not with the root of A.

### 4. Generating multiple-sequence alignments for BEAST

To generate a multiple sequence alignmentℳ = (*S*_1_, …, *S*_*n*_) coherent with an allelic tree *A*, we begin by associating to the root of *A* a sequence *S*_0_ of length greater than or equal to the number of mutations in A, composed strictly of the thymine DNA base (T). The sequence *S*_0_ is then propagated through the branches of *A*. If we index the nodes of *A* by *i*, where *i* = 0 is the root, *i*^′^ is a child of *i* and *b*_*i,i*_ ′ is the length of the branch (i.e. number of mutations) connecting node *i*^′^ to node *i*, then the sequence *S*_*i*_′ is obtained from the sequence *S*_*i*_ by drawing *b*_*i*_′, *i* positions in *S*_*i*_ where the nucleotide is *T* without replacement, and mutating the nucleotides at those positions from *T* to *G*. The length of the sequences at the leaves of *A* is by construction equal to the length of *S*_0_. In addition to providing BEAST with a multiple sequence alignment consistent with *A*, we specify priors on all the different subsets of sequences which have a common ancestor in *A*. This ensures that all the trees sampled during the inference performed by BEAST are constrained to have the same topology as *A*.

To make as fair a comparison as possible between our inference model and BEAST, we ensure that the models specified in BEAST are consistent with the parameters used while generating allelic trees. The generation process of mutations ensures that each site in ℳ mutates with equal probability equal to the inverse of the length of a sequence, either once or not at all (mutation sites are picked without replacement). Since we assume all sites on the sequence are homogeneous, specifying a rate model among sites becomes unnecessary. For the mutation model, we therefore pick a simple Jukes and Cantor 1969 model [54] with no gamma rate variation. The tree model we chose is a simple birth-death model with rates *λ* and *δ* drawn from a prior specified in the main text.

**FIG S4:**
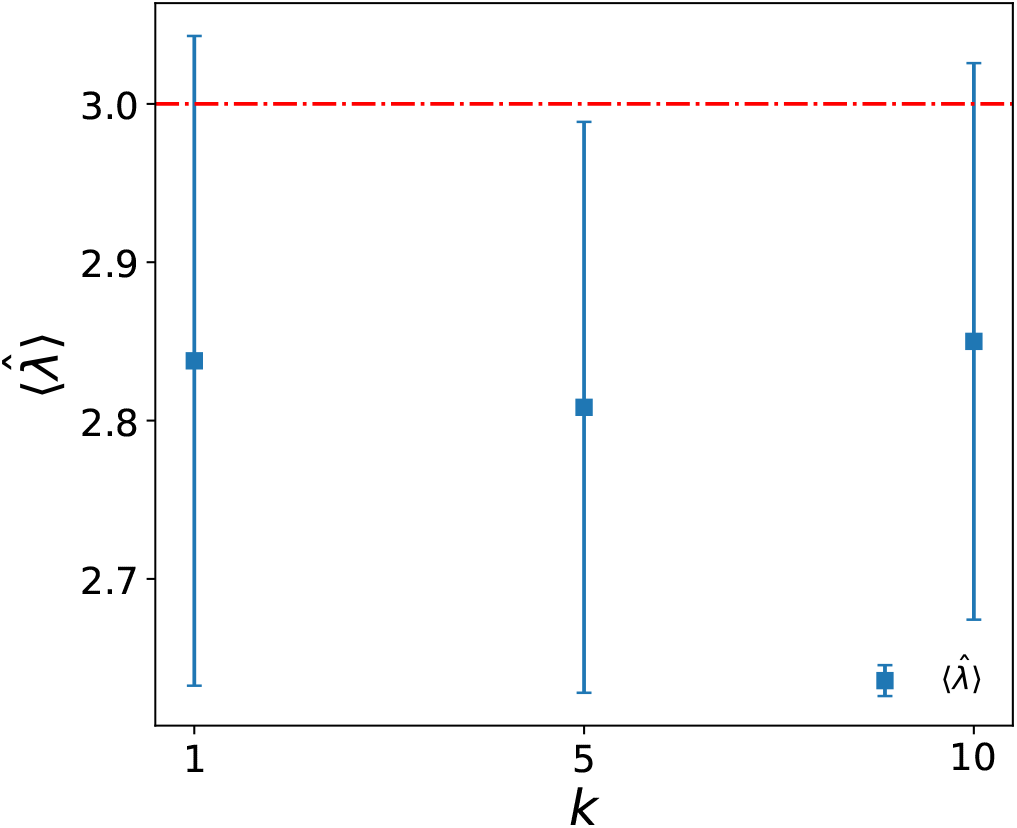
Dependence of 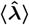 on *k*. Average 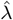 inferred as a function of the number *k* of backbones sampled at every step of the expectation maximization algorithm for *λ*^∗^ = 3, *δ*^∗^ = 1, *µ*^∗^ = 1, *ρ*^∗^ = 1, *T* = 3 and *N* = 1. Numerical experiments are repeated 100 times. Increasing *k* leads to a slight decrease in variance, however no improvement in 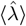is observed. Error bars are given by the standard deviation of multiple inferences

**FIG S5:**
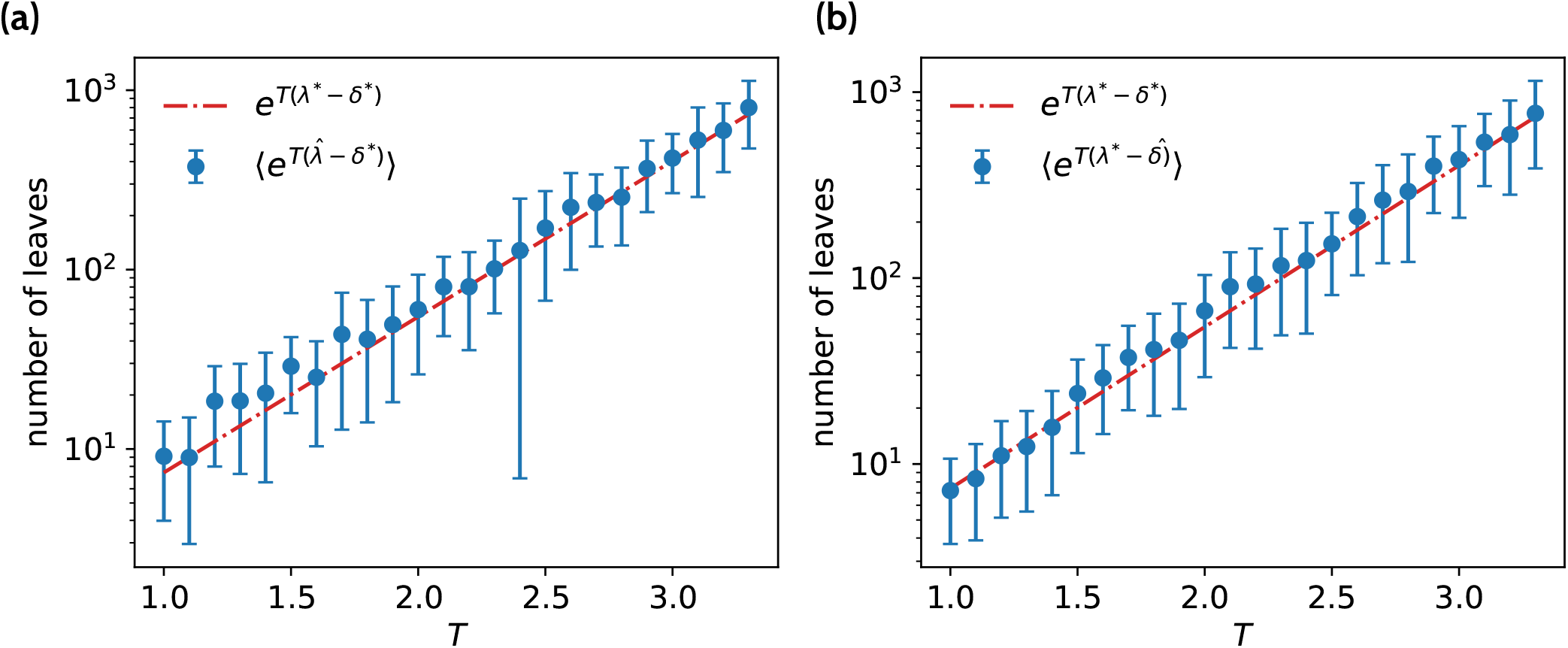
Estimated average number of leaves in *C*. (a,b) Average inferred number of leaves in *C* calculated by averaging over 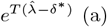 and 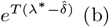 while varying *T*, for *λ*^∗^ = 3, *δ*^∗^ = 1, *µ*^∗^ = 1, *ρ*^∗^ = 1 and *N* = 1. No bias is present in the estimation in the low data regime. Numerical experiments are repeated 100 times. Error bars are given by the standard deviation of multiple inferences.

## References

[1] Jones MG, Yang D, Weissman JS (2023) New tools for lineage tracing in cancer in vivo. Annual Review of Cancer Biology 7:111–129.

[2] Yang D, et al. (2022) Lineage tracing reveals the phylodynamics, plasticity, and paths of tumor evolution. Cell 185:1905–1923.e25.

[3] Simeonov KP, et al. (2021) Single-cell lineage tracing of metastatic cancer reveals selection of hybrid emt states. Cancer Cell 39:1150–1162.e9.

[4] DeWitt, William S I, Mesin L, Victora GD, Minin VN, Matsen, Frederick A I (2018) Using Genotype Abundance to Improve Phylogenetic Inference. Molecular Biology and Evolution 35:1253–1265.

[5] Turner JS, et al. (2021) SARS-CoV-2 mRNA vaccines induce persistent human germinal centre responses. Nature 596:109–113.

[6] Hoehn KB, et al. (2021) Human B cell lineages associated with germinal centers following influenza vaccination are measurably evolving. Elife 10.

[7] Uduman M, et al. (2011) Detecting selection in immunoglobulin sequences. Nucleic Acids Res. 39:W499– 504.

[8] Yaari G, Benichou JIC, Vander Heiden JA, Kleinstein SH, Louzoun Y (2015) The mutation patterns in b-cell immunoglobulin receptors reflect the influence of selection acting at multiple time-scales. Philos. Trans. R. Soc. Lond. B Biol. Sci. 370:20140242.

[9] Nourmohammad A, Otwinowski J, Luksza M, Mora T, Walczak AM (2019) Fierce selection and interference in b-cell repertoire response to chronic HIV-1. Mol. Biol. Evol. 36:2184–2194.

[10] Horns F, Vollmers C, Dekker CL, Quake SR (2019) Signatures of selection in the human antibody repertoire: Selective sweeps, competing subclones, and neutral drift. Proc. Natl. Acad. Sci. U. S. A. 116:1261–1266.

[11] Perié L, Duffy KR, Kok L, de Boer RJ, Schumacher TN (2015) The branching point in erythro-myeloid differentiation. Cell 163:1655–1662.

[12] Lee-Six H, et al. (2018) Population dynamics of normal human blood inferred from somatic mutations. Nature 561:473–478.

[13] Fabre MA, et al. (2022) The longitudinal dynamics and natural history of clonal haematopoiesis. Nature 606:335–342.

[14] Mitchell E, et al. (2022) Clonal dynamics of haematopoiesis across the human lifespan. Nature 606:343–350.

[15] Stadler T, Pybus OG, Stumpf MPH (2021) Phylodynamics for cell biologists. Science 371:eaah6266.

[16] Spencer Chapman M, et al. (2021) Lineage tracing of human development through somatic mutations. Nature 595:85–90.

[17] Park S, et al. (2021) Clonal dynamics in early human embryogenesis inferred from somatic mutation. Nature 597:393–397.

[18] Coorens THH, et al. (2021) Extensive phylogenies of human development inferred from somatic mutations. Nature 597:387–392.

[19] Martincorena I, et al. (2017) Universal patterns of selection in cancer and somatic tissues. Cell 171:1029– 1041.e21.

[20] Bouckaert R, et al. (2014) Beast 2: A software platform for bayesian evolutionary analysis. PLOS Computational Biology 10:1–6.

[21] Karcher MD, Palacios JA, Lan S, Minin VN (2017) phylodyn: an R package for phylodynamic simulation and inference. Mol. Ecol. Resour. 17:96–100.

[22] Johnson B, Shuai Y, Schweinsberg J, Curtius K (2023) clonerate: fast estimation of single-cell clonal dynamics using coalescent theory. Bioinformatics 39.

[23] Sagulenko P, Puller V, Neher RA (2018) TreeTime: Maximum-likelihood phylodynamic analysis. Virus Evol. 4.

[24] Neher RA, Russell CA, Shraiman BI (2014) Predicting evolution from the shape of genealogical trees. eLife 3:e03568.

[25] Dieselhorst T, Berg J (2024) Phylodynamic inference on single-cell data.

[26] Grenfell BT, et al. (2004) Unifying the epidemiological and evolutionary dynamics of pathogens. Science 303:327–332.

[27] Gao Y, Feder AF (2024) Detecting branching rate heterogeneity in multifurcating trees with applications in lineage tracing data. bioRxiv.

[28] Zwaans A, Seidel S, Manceau M, Stadler T (2024) Bayesian phylodynamics of early vertebrate development in BEAST 2.

[29] Prillo S, Ravoor A, Yosef N, Song YS (2023) ConvexML: Scalable and accurate inference of single-cell chronograms from CRISPR/Cas9 lineage tracing data.

[30] Feng J, et al. (2021) Estimation of cell lineage trees by maximum-likelihood phylogenetics. The Annals of Applied Statistics 15:343 – 362.

[31] Seidel S, Stadler T (2022) TiDeTree: a bayesian phylogenetic framework to estimate single-cell trees and population dynamic parameters from genetic lineage tracing data. Proc. Biol. Sci. 289:20221844.

[32] Lambert A, Stadler T (2013) Birth–death models and coalescent point processes: The shape and probability of reconstructed phylogenies. Theoretical Population Biology 90:113–128.

[33] Nee S, May RM, Harvey PH (1994) The reconstructed evolutionary process. Philos. Trans. R. Soc. Lond. B Biol. Sci. 344:305–311.

[34] Morlon H, Parsons TL, Plotkin JB (2011) Reconciling molecular phylogenies with the fossil record. Proc. Natl. Acad. Sci. U. S. A. 108:16327–16332.

[35] Minh BQ, et al. (2020) Iq-tree 2: New models and efficient methods for phylogenetic inference in the genomic era. Molecular Biology and Evolution 37:1530–1534.

[36] Stamatakis A (2014) Raxml version 8: a tool for phylogenetic analysis and post-analysis of large phylogenies. Bioinformatics 30:1312–1313.

[37] Spisak N, de Manuel M, Milligan W, Sella G, Przeworski M (2024) The clock-like accumulation of germline and somatic mutations can arise from the interplay of DNA damage and repair. PLoS Biol. 22:e3002678.

[38] Abascal F, et al. (2021) Somatic mutation landscapes at single-molecule resolution. Nature 593:405–410.

[39] Victora GD, Nussenzweig MC (2022) Germinal centers. Annu. Rev. Immunol. 40:413–442.

[40] Pae J, et al. (2021) Cyclin D3 drives inertial cell cycling in dark zone germinal center B cells. The Journal of Experimental Medicine 218:e20201699.

[41] Williams N, et al. (2022) Life histories of myeloproliferative neoplasms inferred from phylogenies. Nature 602:162–168.

[42] Van Egeren D, et al. (2021) Reconstructing the lineage histories and differentiation trajectories of individual cancer cells in myeloproliferative neoplasms. Cell Stem Cell 28:514–523.e9.

[43] Schwartz R, Schäffer AA (2017) The evolution of tumour phylogenetics: principles and practice. Nat. Rev. Genet. 18:213–229.

[44] McKenna A, et al. (2016) Whole-organism lineage tracing by combinatorial and cumulative genome editing. Science 353:aaf7907.

[45] Lupo C, Spisak N, Walczak AM, Mora T (2022) Learning the statistics and landscape of somatic mutation-induced insertions and deletions in antibodies. PLoS Comput. Biol. 18:e1010167.

[46] Kreer C, et al. (2023) Probabilities of developing HIV1 bNAb sequence features in uninfected and chronically infected individuals. Nat. Commun. 14:7137.

[47] Cobey S, Wilson P, Matsen, 4th FA (2015) The evolution within us. Philos. Trans. R. Soc. Lond. B Biol. Sci. 370:20140235.

[48] Marchi J, Lässig M, Walczak AM, Mora T (2021) Antigenic waves of virus-immune coevolution. Proc. Natl. Acad. Sci. U. S. A. 118:e2103398118.

[49] Briney B, Inderbitzin A, Joyce C, Burton DR (2019) Commonality despite exceptional diversity in the baseline human antibody repertoire. Nature 566:393–397.

[50] Barido-Sottani J, Vaughan TG, Stadler T (2020) A multitype birth-death model for bayesian inference of lineage-specific birth and death rates. Syst. Biol. 69:973– 986.

[51] Victora GD, et al. (2010) Germinal center dynamics revealed by multiphoton microscopy with a photoactivatable fluorescent reporter. Cell 143:592–605.

[52] Louca S, Pennell MW (2020) Extant timetrees are consistent with a myriad of diversification histories. Nature 580:502–505.

[53] Werner B, et al. (2020) Measuring single cell divisions in human tissues from multi-region sequencing data. Nat. Commun. 11:1035.

[54] Jukes TH, Cantor CR (1969) in Mammalian Protein Metabolism (Elsevier), pp 21–132.

